# Early life stress exposure increases susceptibility to high fat/high sucrose diet in female mice

**DOI:** 10.1101/2022.07.14.500119

**Authors:** Jenna M. Frick, Olivia C. Eller, Rebecca M. Foright, Brittni M. Levasseur, Xiaofang Yang, Ruipeng Wang, Michelle K. Winter, Maura F. O’Neil, E. Matthew Morris, John P. Thyfault, Julie A. Christianson

## Abstract

Exposure to stress early in life has been associated with adult-onset co-morbidities such as chronic pain, metabolic dysregulation, obesity, and inactivity. We have established an early life stress model using neonatal maternal separation (NMS) in mice, which displays evidence of increased body weight and adiposity, widespread mechanical allodynia, and hypothalamic-pituitary-adrenal axis dysregulation in male mice. Early life stress and consumption of a western style diet contribute to the development of obesity, however, relatively few pre-clinical studies have been performed in female rodents, which are known to be protected against diet induced obesity and metabolic dysfunction. In this study we gave naïve and NMS female mice access to a high-fat/high-sucrose (HFS) diet beginning at 4 weeks of age. Robust increases in body weight and fat were observed in HFS-fed NMS mice during the first 10 weeks on the diet, driven partly by increased food intake. Female NMS mice on a HFS diet showed widespread mechanical hypersensitivity compared to either naïve mice on a HFS diet or NMS mice on a control diet. HFS diet-fed NMS mice also had impaired glucose tolerance and fasting hyperinsulinemia. Strikingly, female NMS mice on a HFS diet showed evidence of hepatic steatosis with increased triglyceride levels and altered glucocorticoid receptor levels and phosphorylation state. They also exhibited increased energy expenditure as observed via indirect calorimetry and expression of pro-inflammatory markers in perigonadal adipose. Altogether, our data suggest that early life stress exposure increased the susceptibility of female mice to develop diet-induced metabolic dysfunction and pain-like behaviors.

## Introduction

Over 40% of adults in the United States have a body mass index that would be classified as obese (1, 2). This high prevalence of obesity not only increases the likelihood of negative health outcomes including cardiovascular disease, cancer, and type 2 diabetes (3–7), but also constitutes a large financial burden, with an estimated yearly cost of 147 billion dollars or nearly 9% of annual medical costs (8). Considering these medical and financial implications, it is imperative that we better understand the physiologic mechanisms and lifestyle factors that contribute to obesity.

Exposure to early life stress (ELS) has been found to result in poor physical and mental health outcomes, such as chronic pain conditions, mood disorders, substance abuse, inactivity, and obesity (9–13). Males and females exposed to physical and sexual abuse early in life are two times more likely to develop severe obesity (14) and have an average body weight nearly nine pounds higher than their un-abused peers (12, 15–17). ELS exposure, which can occur in many forms including neglect, abuse, and premature birth, is highly prevalent in the U.S. as over 64% of the adult population reports at least one adverse childhood experience (9). The Child Maltreatment Report estimates that 678,000 or 9 out of every 1000 children were victims of abuse or neglect in 2018 with the highest rate of maltreatment among those under the age of one (18). Exposure to negative environmental stimuli during early development, which consists of a period of high neural plasticity, can have long-term detrimental consequences (18–23).

In addition to ELS, other lifestyle factors may also be contributing to poor metabolic outcomes. Consumption of a diet high in fat and sucrose content has been consistently associated with weight gain and impaired glucose homeostasis (24–27). This type of diet is highly prevalent in the U.S. due to its palatability, affordability, and accessibility (1, 2, 28). Importantly, high-fat/high-sucrose (HFS) diets are often consumed by low socioeconomic status (SES) populations that are also more prone to ELS, particularly in the form of food insecurity, poor housing conditions, and unstable home life (29, 30). It is therefore very likely that individuals exposed to ELS may also be consuming a HFS diet, and the combination of these factors may result in worsened metabolic health.

Rodent models are often used to examine the underlying mechanisms contributing to weight gain. Although female rodents show some resistance to obesity due to the protective effects of estrogen (31, 32), a large proportion of premenopausal female women are documented as obese or overweight (33). It is likely that a combination of biological and lifestyle factors are required to overcome the protective effects of estrogen. To determine how ELS and HFS diet impact metabolism in a female rodent, we performed neonatal maternal separation (NMS) and weaned female pups onto either control chow or a HFS diet. Measures of body weight and fat gain, glucose tolerance, mechanical sensitivity, hepatic steatosis, indirect calorimetry, and inflammatory markers in adipose were carried out either over time or after 17-25 weeks on the diet.

## Methods

### Animals

All experiments in this study were performed on female C57BI/6 mice (Charles River, Wilmington, MA) born and housed in the Laboratory Animal Resources Facility at the University of Kansas Medical Center. All mice received food and water ad libitum and were housed on a 12-hour light cycle (600 to 1800 h). All research conformed to the National Institutes of Health Guide for the Care and Use of Laboratory Animals and was approved by the University of Kansas Medical Center Institutional Animal Care and Use Committee (protocol number 2019-2501).

### Experimental Design

Our experimental design is schematized in Fig. 1. Mice underwent neonatal maternal separation (NMS) or remained unhandled during the first 3 weeks of life. Mice were introduced to a high-fat/high-sucrose (HFS) diet or control diet at 4 weeks of age and remained on their assigned diet throughout the duration of the study. After 20-25 weeks on the diet mice were housed in indirect calorimetry cages for 7 days prior to euthanasia. Group sizes are reported in the figure legends and were based on prior studies from our laboratories.

**Figure 1.**
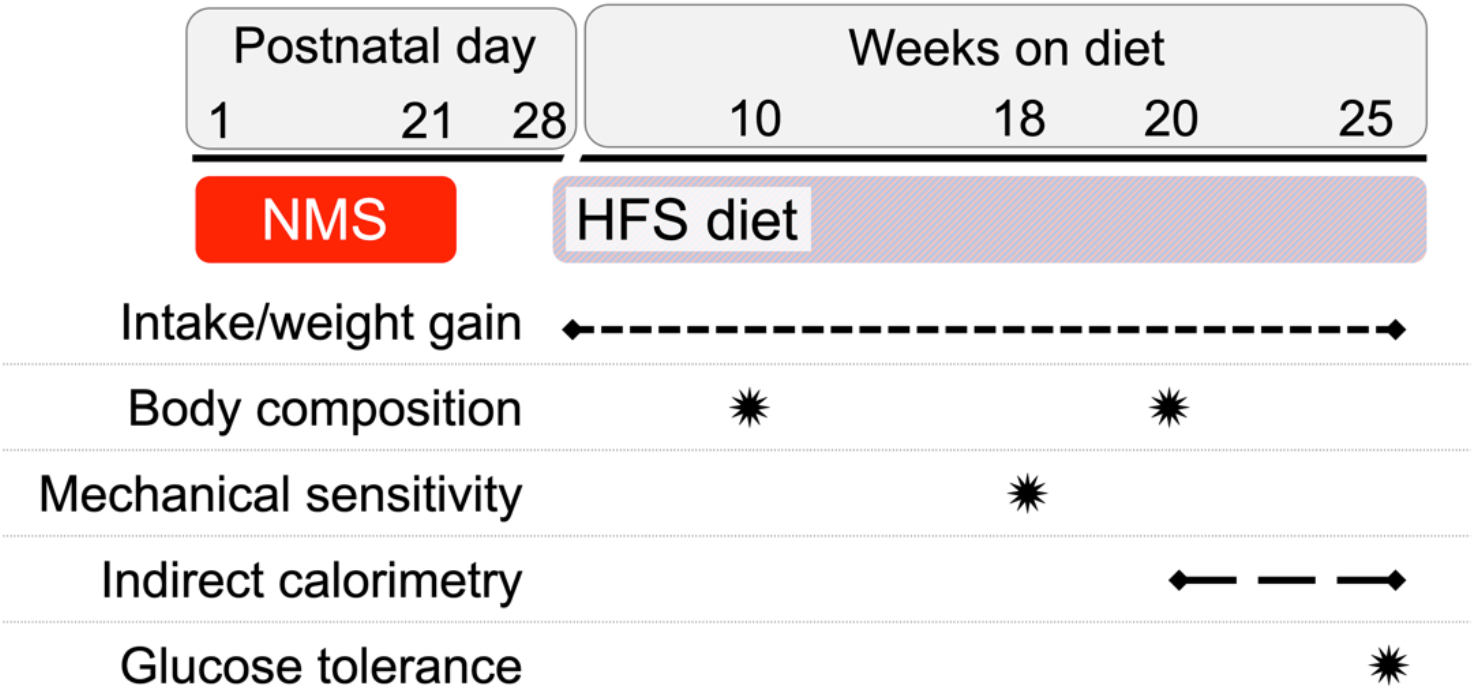
Schematic of the experimental approach. Female mice underwent NMS from P1 to P21, or remained unhandled (naïve), and were weaned and pair-housed with littermates on P22. On P28, half of the cages were given ad libitum access to a HFS diet and half remained on a control diet. Weight gain and diet consumed were measured on a weekly basis. Body composition analysis occurred after 10- and 20-weeks on the diet. Mechanical withdrawal thresholds were measured from the hindpaw and perigenital region after 18 weeks on the diet. Indirect calorimetry was performed for one week between 20-25 weeks on the diet. End point measurements including fasting insulin and glucose, glucose tolerance, and adipose and liver analyses were taken after 29-weeks on the diet.

### Neonatal Maternal Separation (NMS)

Pregnant dams were delivered to the animal facility during the last week of gestation. Beginning on postnatal day 1 (P1) until P21, NMS litters were removed and placed *en masse* into clean glass beakers, containing small amounts of their home cage bedding material, and held at 33°C and 50% humidity for 180 min (11am – 2pm) daily (34). Naïve mice remained undisturbed in their home cage without handling outside of normal husbandry care. No significant difference in number of pups per litter was observed between naïve (7.5 ± 0.5) and NMS (7 ± 0.6) litters. All mice were weaned on P22 and caged with same-sex littermates. At 4 weeks of age mice were pair-housed with a littermate to eliminate potential stress associated with single housing. Although only female mice are reported here, all pups in each litter (male and female) underwent NMS, or remained unhandled, until weaning.

### High-fat/high-sucrose diet

At 4 weeks of age, half of the naïve and NMS mice were transitioned to a HFS diet (20% kcal protein, 35% kcal carbohydrate (17% sucrose), 45% kcal fat; 4.7 kcal/g; Research Diets, Inc. New Brunswick, NJ; cat. D12451) or remained on a control diet (20% kcal protein, 70% kcal carbohydrate (3.5% sucrose), 10% kcal fat; 3.8 kcal/g; Research Diets, cat. D12110704).

### Food consumption and feed efficiency

Body weight and food consumption per pair of mice were monitored weekly. Feed efficiency was quantified every week using the following equation: ((weight change per pair)/(calories consumed per pair))*1000. These data are presented as average caloric intake and feed efficiency over weeks 1-10 and 11-20 that the mice were on HFS or control diet.

### Body Composition

Mice were weighed weekly and body composition was measured via qMRI using an EchoMRI 1100 (EchoMRI LLC, Houston, TX). Total weight, percent body fat, and lean mass were quantified at 10 and 20 weeks on the diet.

### Von Frey Monofilament Testing

At 17 weeks on the diet, hindpaw and perigenital mechanical sensitivity were measured using von Frey monofilaments as previously described (34). Mice were acclimated to a soundproof testing room in their home cages for 30 minutes prior to testing. Mice were placed inside of individual clear plastic chambers (11 cm x 5 cm x 3.5 cm) on a wire mesh screen elevated 55 cm above a table. The up-down method was performed to test mechanical sensitivity using a standard set of von Frey monofilaments (1.65, 2.36, 2.83, 3.22, 3.61, 4.08, 4.31, 4.74 g, Stoelting Co., Wood Dale, IL, United States) (35). Beginning with the 3.22 g monofilament, mice received a single application to the perigenital or hindpaw region and a positive response (considered a jump, jerk, or licking of the affected area) was followed by the next smaller filament. The experimenter continued to move up or down the series depending on the previously elicited response, for an additional four applications after the first positive response was observed for a minimum of five or a maximum of nine total monofilament applications. The value in log units of the final von Frey monofilament applied in the trial series was used to calculate a 50% g threshold for each mouse and group means were determined as previously described (36).

### Indirect calorimetry cages

An 8-channel Promethion (Sable Systems International, Las Vegas, NV, USA) system was used for indirect calorimetry. The Promethion GA-3 gas analyzer measured H2O, CO2, and O2 levels in each cage and the airflow through each cage was held at 2,000 mL/min. Cages were held in a separate room that was matched for the temperature and light cycle of the home room. All mice were acclimated to single-caging conditions for two weeks prior to housing in the indirect calorimetry cages. Mice were individually housed in Promethion indirect calorimetry cages for seven days total. The first four days were used for acclimation to the system and data from the last three days in the cages were analyzed via Sable Systems International EXPEDATA Data Acquisition & Analysis Release 1.9.27. Data were binned per 24-hour period and values for the three days were averaged and extrapolated to weekly values.

### Glucose tolerance test

After 20-25 weeks on the diet, mice were subject to a glucose tolerance test. Following a 6-hour fast, mice were given an intraperitoneal injection of glucose at 1 g/kg body weight. Blood glucose levels were measured via tail clip immediately before glucose injection and 15, 30, 60, and 120 minutes following the injection using a colorimetric assay (PGO enzyme preparation and dianisidine dihydrochloride, Sigma-Aldrich, St. Louis, MO). Area under the curve measurements were calculated using the trapezoidal method (37).

### Fasting insulin level

The initial fasting blood collection described above was also used to measure serum insulin levels using an insulin ELISA kit according to the manufacturer’s instructions (ALPCO, Salem, NH). HOMA-IR was calculated using the formula: Fasting plasma glucose (mmol/L) X fasting serum insulin (μU/mL) / 22.5.

### Histology

During dissection, a portion of the periovarian adipose tissue and the liver were fixed in 4% paraformaldehyde for 48 hours prior to storage in 70% ethanol. Fixed tissues were embedded in paraffin wax, sectioned, and mounted on glass slides for hematoxylin and eosin (H&E) staining. Bright field images were captured at 10x optical magnification with a Nikon Eclipse 90i upright microscope (Nikon, Melville, NY). Adipocyte area was measured using ImageJ in a semiautomated method as described by Parlee in 2014 (38). H&E stained liver sections were examined via bright field to determine a non-alcoholic fatty liver disease activity (NAS) score. Liver sections were examined to determine the level of steatosis, lobular inflammation, and hepatocyte ballooning, and the sum of these scores was calculated as the total NAS score (39).

### Liver Triglyceride Assay

Frozen liver tissue was manually crushed and 30 mg per sample was homogenized in 1mL of 2:1 chloroform:methanol via a TissueLyser II (Qiagen). Samples were stored at 4 °C overnight with agitation. 1 mL of 4mM MgCl2 was added to each sample followed by centrifugation for 1 hour at 3100 rpm. 500 μL of organic phase was transferred to a new tube and tubes were left open to evaporate overnight. Dried lipids were reconstituted in 100 μL of 3:2 butanol:triton-x114 mix. A glycerol standard curve (Sigma, Saint Louis, MO) was used at 2.5, 1.25, 0.625, and 0.3125 mg/mL. 3μL of samples and standards were loaded into a 96 well plate and the solvent was allowed to evaporate for 5 minutes. 300 μL of 4:1 Free Glycerol Reagent:Triglyceride Reagent (Sigma) was added to each well and the plate was incubated at 37 °C for 15 minutes. The plate was read at 540 nm and samples were compared to the glycerol standard. Values were converted to nMol/g of tissue in wet weight for final concentration.

### Liver Western blot

Total protein was isolated from the liver using Cell Extraction Buffer containing Halt protease and phosphatase inhibitors (ThermoFisher Scientific, Waltham, MA) and Na3VO4. A Dc protein assay (ThermoFisher Scientific) was used to determine protein concentration and 400μg samples were reduced, subjected to SDS-PAGE (Criterion 4% to 12% Bis-Tris gels; Bio-Rad), and transferred to Nitrocellulose membrane (Whatman GmbH, Dassel, Germany). Nonspecific binding was blocked by incubation in 5% milk in Tris-buffered saline with Tween-20 (TBST) for 1h. Membranes were then incubated overnight at 4°C with primary antisera to GR, pSer211-GR, or GAPDH (1:1000, Cell Signaling, Danvers, MA). Membranes were then washed with TBST and incubated with anti-rabbit secondary antibody (1:10,000; Cell Signaling Technology) for 1 h. Densitometry was performed using Quantity One 4.6.9 software (Bio-Rad).

### RT-PCR

Mice were overdosed with inhaled isoflurane (>5%). Retroperitoneal and periovarian adipose tissue were dissected, weighed, and immediately frozen in liquid nitrogen and stored at −80 °C. Frozen periovarian fat was manually crushed and then homogenized via a TissueLyser II (Qiagen, Valencia, CA). mRNA was extracted from 0.100 grams of crushed tissue via RNeasy Lipid Tissue Mini Kit (Qiagen, Valencia, CA). mRNA was checked for concentration and purity using NanoDrop 2000 (Thermo Fisher Scientific, Wilmington, DE). mRNA was synthesized into cDNA using the iScript cDNA Synthesis Kit (Bio-Rad, Hercules, CA), BIORAD T100 Thermal Cycler, and stored at −20 °C. Quantitative RT-PCR was performed using SsoAdvanced SYBR Green Supermix (Bio-Rad) and a Bio-Rad iCycler IQ real time PCR system with indicated 20 μM primers (Table 1; Integrated DNA Technologies, Coralville, IA). Samples were run in triplicate and negative control reactions were run with each amplification series. To reduce variability among efficiency due to fluctuations in baseline fluorescence, the raw PCR data was imported to the LinRegPCR software (v2012.3) (40) and PCR efficiency values were derived for each individual sample. Threshold cycle values were subtracted from the housekeeping gene PPiB and the fold change over naïve-Control was calculated using the Pfaffl method (41).

**Table 1.**
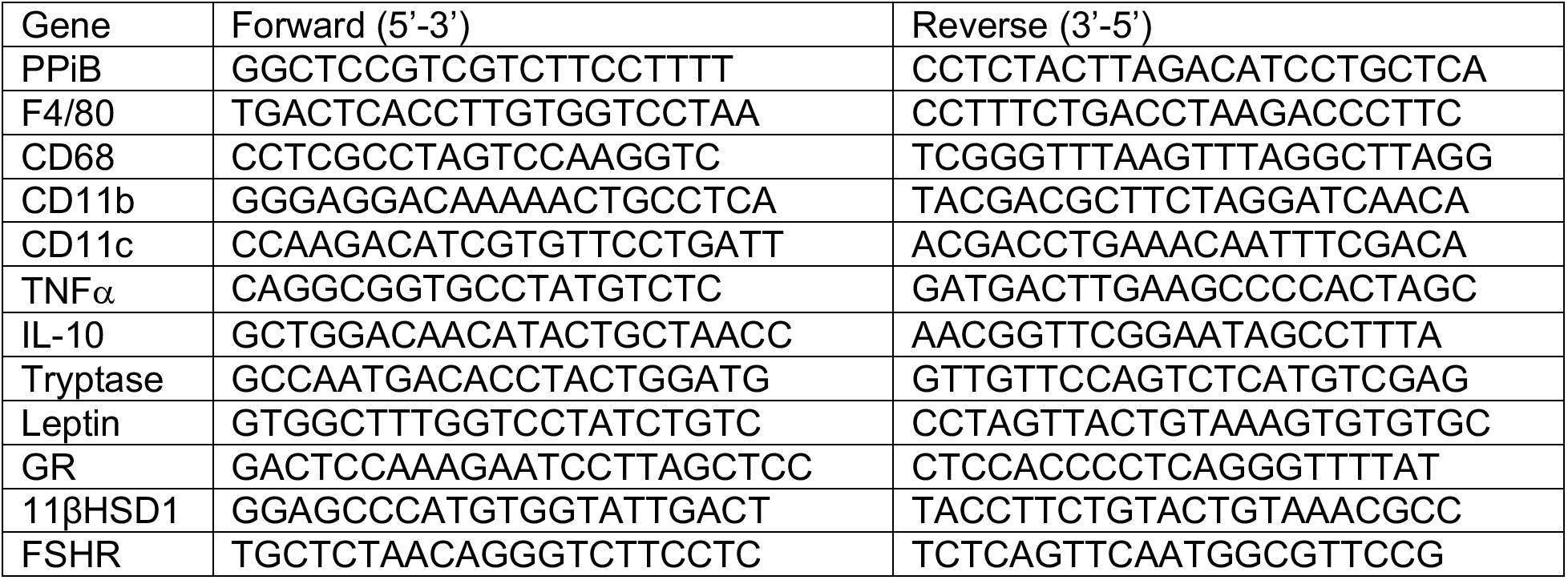
Forward and reverse primers used for RT-PCR analysis of perigonadal adipose tissue.

### Statistics

Calculations of the above measurements were made in Excel (Microsoft, Redmond, WA) and statistical analyses were performed using GraphPad Prism 9 (GraphPad, La Jolla, CA) or IBM SPSS Statistics 27 (IBM Corporation, Armonk, NY). Differences between groups were determined by 2-way ANOVA, with or without repeated measures (RM), and Fisher’s LSD or Bonferroni posttest. ANCOVA was performed in SPSS using weight as the covariate. Statistical significance was set at *p*<0.05. Data are displayed as individual values, where possible, plus mean ± standard error of the mean (SEM).

## Results

### NMS increased rate of weight gain

To observe the effects of NMS and HFS diet consumption on overall health, body weight was measured weekly and body composition was assessed after 10 and 20 weeks on the diet. After only 1 week, NMS mice on HFS diet were significantly heavier than control diet-fed NMS mice and remained so for the duration of the study (Figure 2A). In comparison, naïve mice were not significantly heavier on the HFS diet until after 5 weeks (Figure 2A). After 10 weeks on HFS diet, a significant overall effect of NMS and diet was observed on percent body fat with naïve-HFS and NMS-HFS mice having significantly higher body fat percentage than their control diet-fed counterparts and NMS-HFS mice having higher body fat compared to naïve-HFS mice (Figure 2B). After 20 weeks, HFS diet continued to significantly increase body fat percentage with both HFS diet-fed groups being significantly higher than their control diet-fed counterparts (Figure 2C). No significant effects of diet or NMS were observed on lean mass at either 10 weeks (Figure 2D) or 20 weeks (Figure 2E). Taken together these results indicate that, as expected, HFS diet feeding led to increased body weight and adiposity and that NMS exposure amplified these outcomes at early timepoints.

**Figure 2:**
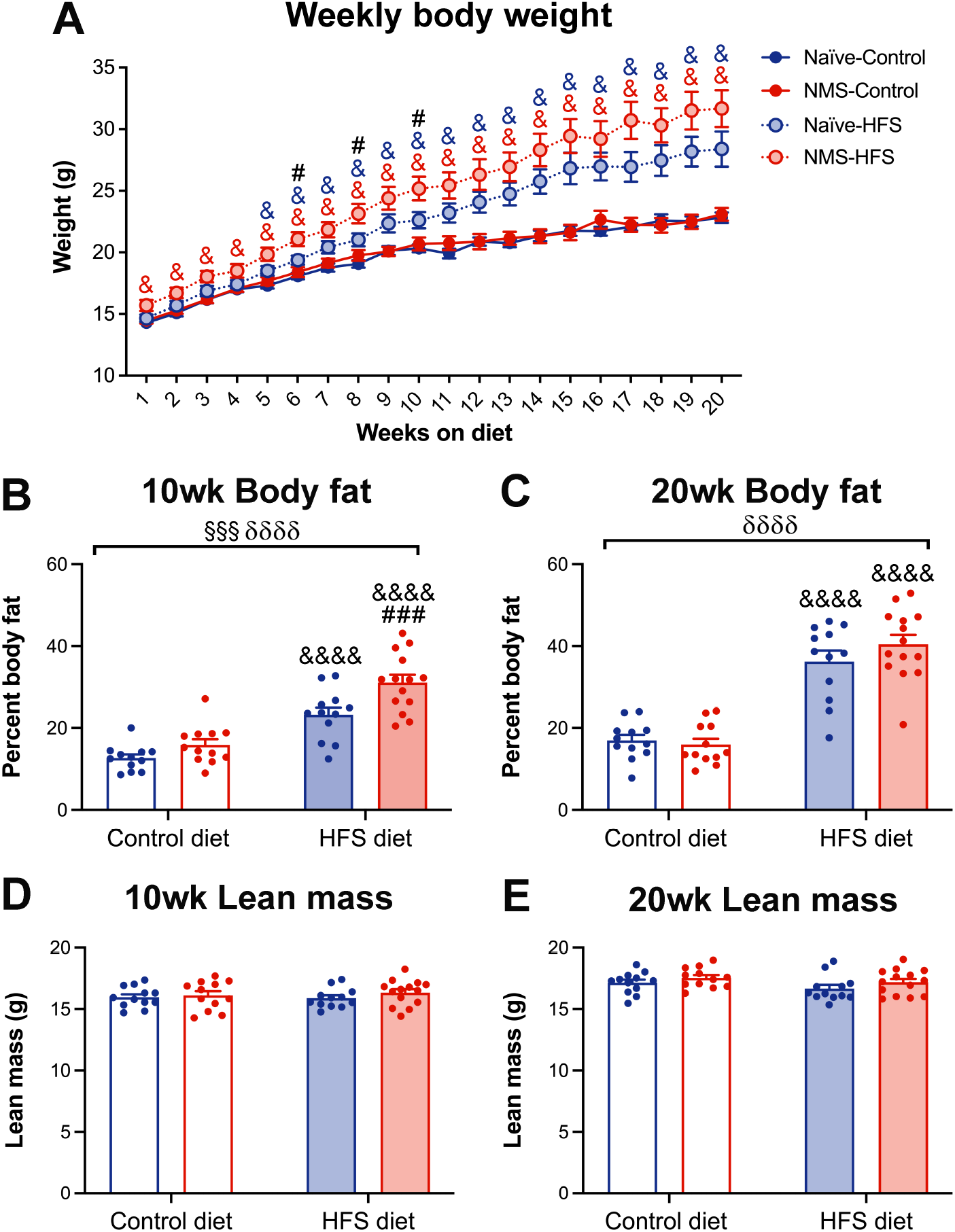
The impact of neonatal maternal separation (NMS) and diet on weekly body weight and body composition after 10 and 20 weeks on high fat/high sucrose (HFS) diet. **A)** A significant impact of diet (*p*<0.0001) and a trend toward an NMS effect (*p*=0.088) was observed on increasing body weight. Significant group differences were observed between naïve-control and naïve-HFS mice (*p*=0.005), NMS-control and NMS-HFS mice (*p*<0.0001), and NMS-HFS and naïve-HFS mice (*p*=0.025). NMS mice on HFS diet were significantly heavier than control diet-fed NMS mice beginning after just 1 week on the diet. In comparison, naïve mice on HFS diet were heavier than control-fed naïve mice beginning after 5 weeks on the diet. NMS-HFS mice were also significantly heavier than naïve-HFS mice at 6, 8, and 10 weeks on the diet. **B)** Body fat percentage after 10 weeks on diet was significantly increased by both NMS (*p*=0.0010) and HFS diet (*p*<0.0001). Both naïve and NMS mice on HFS diet had significantly greater body fat percentage than their control diet-fed counterparts and NMS-HFS mice were significantly higher than naïve-HFS mice. **C)** Body fat percentage after 20 weeks on diet was significantly increased by HFS diet (*p*<0.0001) with no significant difference between naïve and NMS mice on HFS diet. Lean mass at 10 **(D)** and 20 weeks **(E)** was not impacted by either NMS or diet. § and δ denote significant effects of NMS and diet, respectively, three-way RM ANOVA (**A**) or two-way ANOVA (**B-D**). &, &&, &&&, &&&& *p*<0.05, 0.01, 0.001, 0.0001 HFS vs. control, # *p*<0.05 NMS-HFS vs. naïve-HFS, Fisher’s LSD posttest. n=12-14 per group.

### Food consumption was influenced by diet and stress

To examine factors contributing to weight gain, energy intake and feed efficiency were monitored throughout the duration of the study. During the first ten weeks on the diet, we observed a significant overall effect of NMS and diet on increasing average energy intake per pair, such that naïve-HFS mice consumed more calories than naïve-control mice, and NMS-HFS mice consumed significantly more calories than either NMS-control or naïve-HFS mice (Figure 3A). While the effect of NMS was not observed during weeks 11-20, a significant effect of diet on food intake remained (Figure 3B). A significant effect of diet was observed on feed efficiency during both the 1-10 week (Figure 3C) and 11-20 week timepoints (Figure 3D) where HFS diet consumption resulted in greater feed efficiency. These results indicate that both HFS diet consumption and NMS exposure led to higher caloric intake, particularly during the first ten weeks of HFS diet feeding, although only diet significantly impacted feed efficiency.

**Figure 3.**
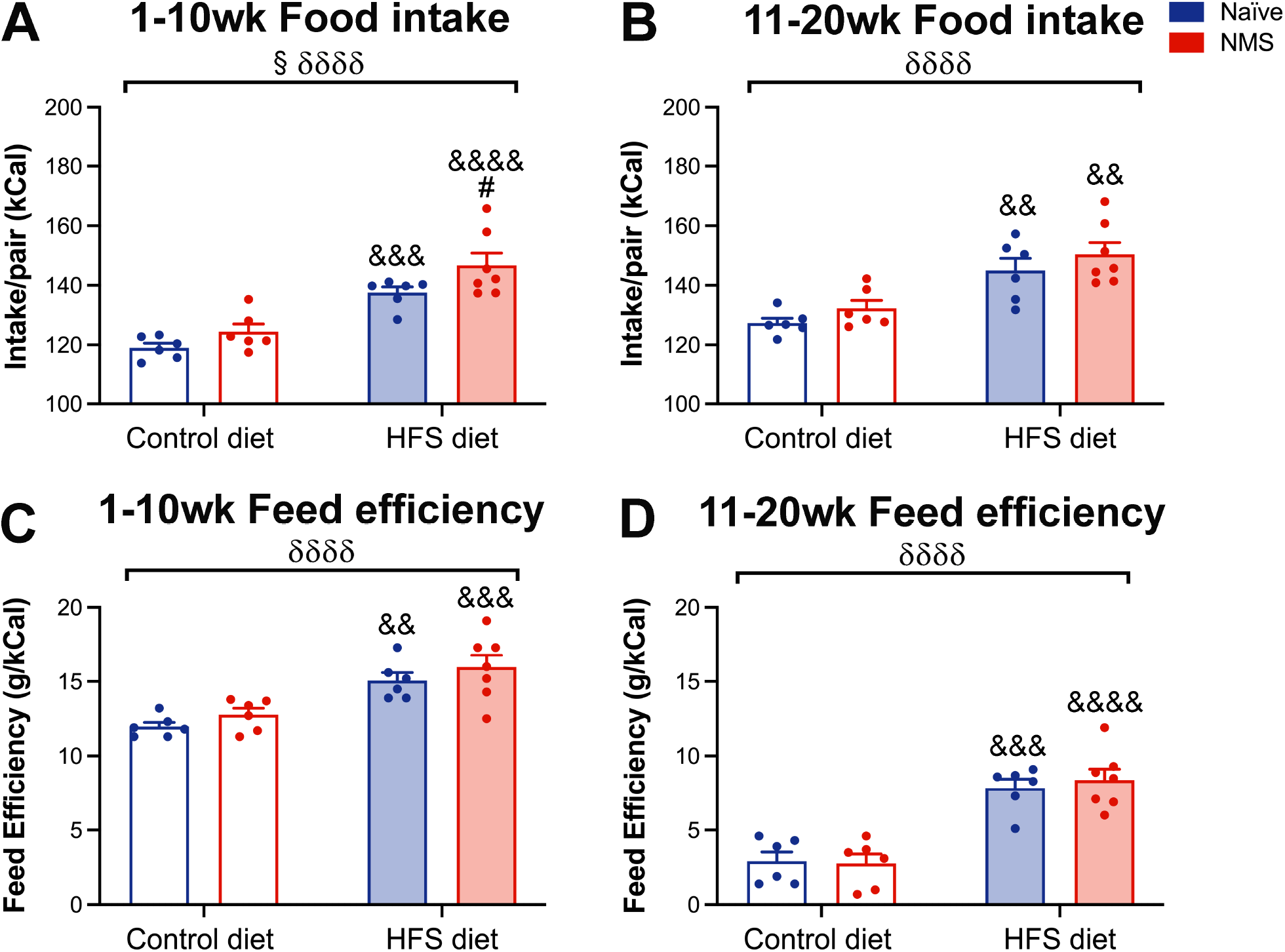
The impact of neonatal maternal separation (NMS) on food intake and feed efficiency was measured weekly and is shown as an average for weeks 1-10 and 11-20 on a high fat/high sucrose (HFS) diet. **A**) Average caloric intake over the first 10 weeks was significantly increased by NMS (*p*=0.0209) and HFS diet (*p*<0.0001), with both HFS groups having higher intake than their control diet-fed counterparts and NMS-HFS having higher intake than naïve-HFS mice. **B**) Average caloric intake during weeks 11-20 remained significantly higher due to HFS diet (*p*<0.0001) with both naïve and NMS mice on HFS diet having higher intake than their control diet-fed counterparts. Feed efficiency was significantly higher due to HFS diet during weeks 1-10 (**C,** *p*<0.0001) and weeks 11-20 (**D,** *p*<0.0001) with both naïve-HFS and NMS-HFS mice having higher feed efficiency than their control diet-fed counterparts. § and δ denote significant effects of NMS and diet, respectively, two-way ANOVA. &&, &&&, &&&& *p*<0.01, 0.001, 0.0001 HFS vs. control, # *p*<0.05 NMS-HFS vs. naïve-HFS, Fisher’s LSD posttest. n=12-14 per group.

### Diet and stress contribute to mechanical allodynia

ELS exposure and high fat diet consumption have both been associated with chronic pain conditions (42–46); accordingly, after 17 weeks on the HFS diet, hindpaw and perigenital mechanical sensitivity were assessed using von Frey monofilaments. Consistent with our previous studies (47–50), hindpaw mechanical withdrawal thresholds were significantly lower due to NMS, HFS diet, and an NMS/HFS diet interaction (Figure 4A). NMS-control and naïve-HFS mice both had significantly lower thresholds than naïve-control mice. Likewise, perigenital mechanical thresholds were significantly lowered due to NMS and HFS diet, with NMS-control and naïve-HFS mice having significantly lower thresholds compared to naïve-control mice (Figure 4B). Hindpaw and perigenital withdrawal thresholds of NMS-HFS mice were also significantly lower than naïve-control mice (*p*<0.0001 and *p*=0.0012, respectively), but were not significantly different from NMS-control or naïve-HFS mice. These results indicate that both NMS exposure and HFS diet resulted in widespread mechanical allodynia.

**Figure 4.**
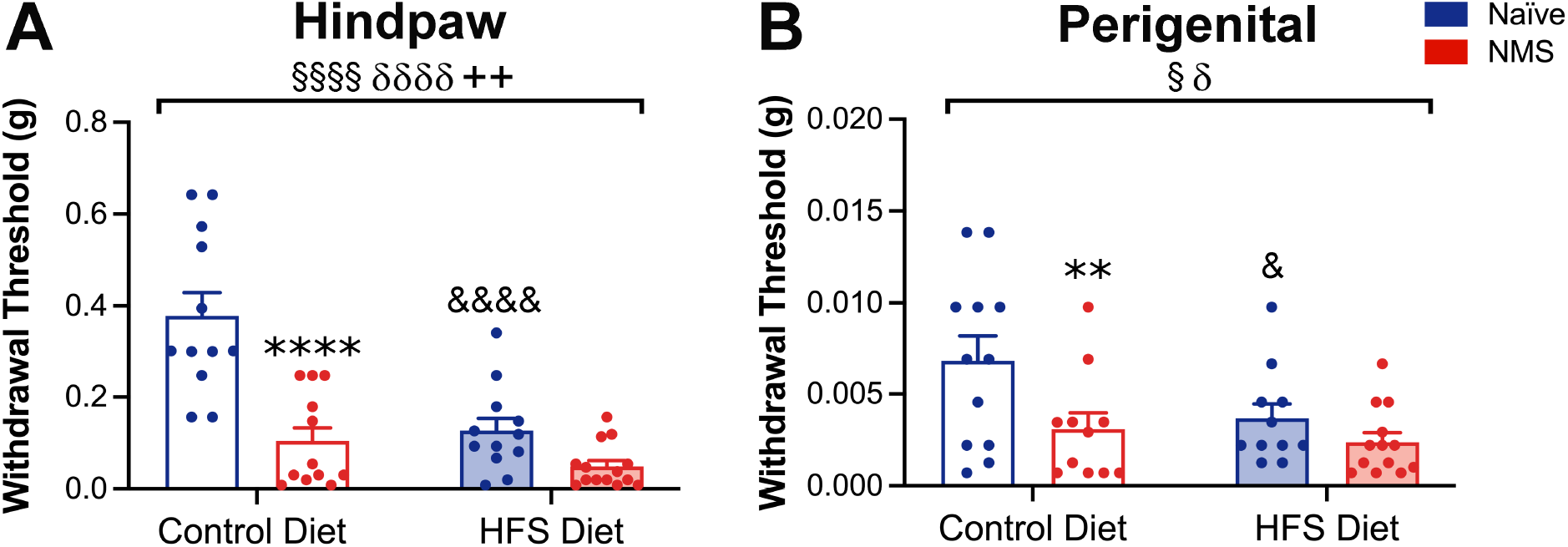
The impact of neonatal maternal separation (NMS) and a high fat/high sucrose (HFS) diet on hindpaw and perigenital mechanical withdrawal threshold was determined after 18 weeks on diet. **A**) A significant effect of NMS (*p*<0.0001), HFS diet (*p*<0.0001), and an interaction effect (*p*=0.0034) was observed on decreasing hindpaw mechanical withdrawal thresholds with NMS-control and naïve-HFS mice having significantly lower thresholds compared to naïve-control mice. **B**) NMS (*p*=0.0101) and HFS diet (*p*=0.0441) significantly lowered perigenital mechanical withdrawal thresholds such that NMS-control and naïve-HFS mice had significantly lower thresholds compared to naïve-control mice. §, δ, and + denote significant effects of NMS, diet, and a NMS/diet interaction respectively, two-way ANOVA. &, &&&& *p*<0.05, 0.0001 HFS vs. control, **, **** *p*<0.01, 0.0001 NMS-control vs. naïve-control, Fisher’s LSD posttest. n=12-14 per group.

### Stress and diet influenced energy expenditure

Energy intake and expenditure were monitored via indirect calorimetry after 20-25 weeks on the diet. Diet and NMS both significantly increased total energy expenditure (TEE), especially in NMS-HFS mice (Table 2). Similarly, resting energy expenditure (REE) was significantly increased by both diet and NMS (Table 2), however, non-resting energy expenditure (NREE) was significantly decreased by diet (Table 2). Energy intake (EI) was also significantly increased by HFS diet (Table 2). Analysis of covariance (ANCOVA) was performed with body weight as the covariate, as body mass is known to influence energy intake and expenditure. The weight adjusted outputs are shown in Table 3. Body weight was determined to be a significant predictor of the variability between groups for both TEE and REE and when body weight was used as a covariate, differences between diets were lost.

**Table 2.**
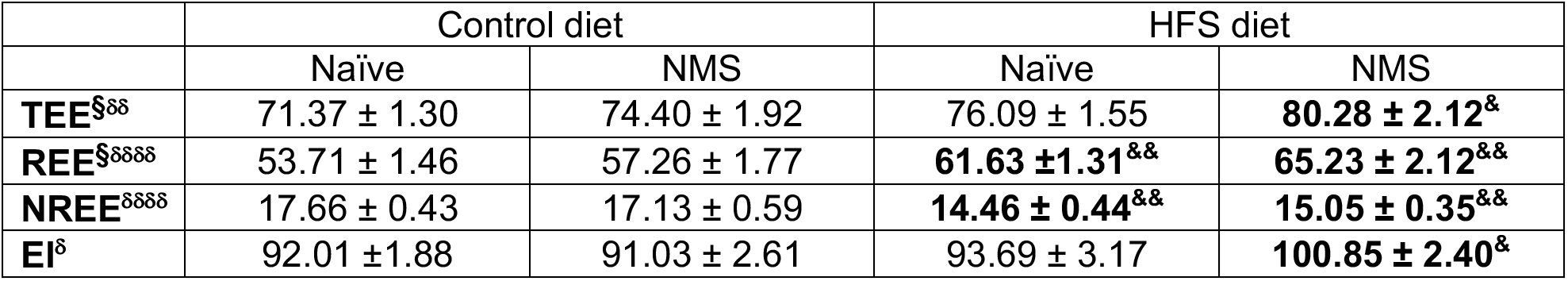
The impact of neonatal maternal separation (NMS) and high fat/high sucrose (HFS) diet on energy intake and expenditure was measured after 20 weeks on the diet. Total energy expenditure (TEE) was significantly increased due to NMS (*p*=0.0469) and diet (*p*=0.0046). Resting energy expenditure (REE) was also significantly increased due to NMS (*p*=0.0419) and diet (*p*<0.0001), while non-resting energy expenditure (NREE) was significantly decreased by diet (*p*<0.0001). Energy intake (EI) was significantly increased due to HFS diet (*p*=0.0311). § and δ denote significant effects of NMS and diet, respectively, and are indicated by bold text, two-way ANOVA. &, && *p*<0.05, 0.01 HFS vs. control, Fisher’s LSD posttest. n=10 per group.

**Table 3.**
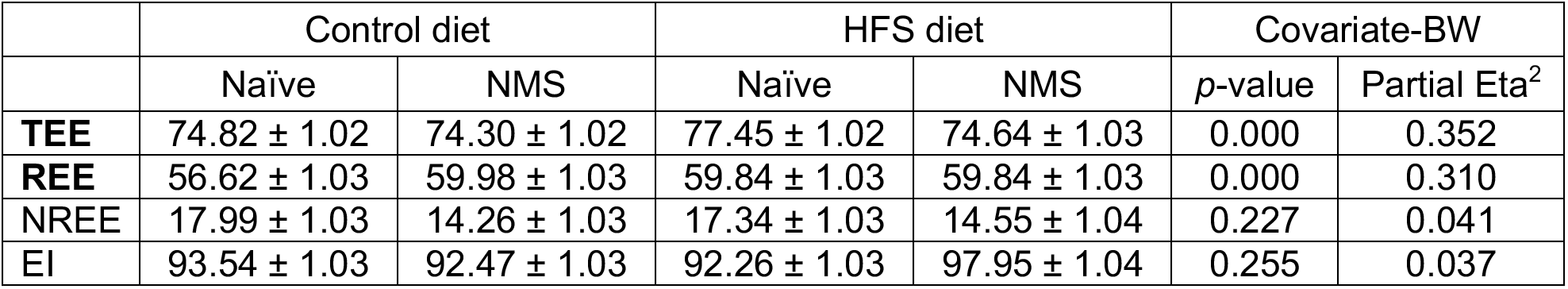
Analysis of covariance (ANCOVA) was performed to determine the impact of body weight (BW) on energy intake and expenditure. A significant effect of body weight was observed on total energy expenditure (TEE, *p*<0.0001) and resting energy expenditure (REE, *p*<0.0001). n=10 per group.

|  | Control diet |  | HFS diet |  | Covariate-BW |  |
| --- | --- | --- | --- | --- | --- | --- |
|  | Naïve | NMS | Naïve | NMS | *p*-value | Partial Eta<sup>2</sup> |
| **TEE** | 74.82 ± 1.02 | 74.30 ± 1.02 | 77.45 ± 1.02 | 74.64 ± 1.03 | 0.000 | 0.352 |
| **REE** | 56.62 ± 1.03 | 59.98 ± 1.03 | 59.84 ± 1.03 | 59.84 ± 1.03 | 0.000 | 0.310 |
| NREE | 17.99 ± 1.03 | 14.26 ± 1.03 | 17.34 ± 1.03 | 14.55 ± 1.04 | 0.227 | 0.041 |
| EI | 93.54 ± 1.03 | 92.47 ± 1.03 | 92.26 ± 1.03 | 97.95 ± 1.04 | 0.255 | 0.037 |

### Stress and HFS diet consumption led to impaired glucose tolerance and insulin sensitivity

Glucose tolerance and fasting serum insulin levels were measured at the end of the study. A significant overall effect of diet and an NMS/diet interaction was observed on glucose tolerance, with NMS-HFS mice having significantly higher serum glucose levels throughout the testing period, compared to NMS-control mice, and at the 120-minute time point compared to naïve-HFS mice (Figure 5A). Similarly, the area under the curve (AUC) calculated from the glucose tolerance test was significantly affected by diet and an NMS/diet interaction (Figure 5B). Both naïve-HFS and NMS-HFS mice had significantly higher serum glucose AUC compared to their naïve counterparts. NMS-HFS mice also had significantly higher glucose AUC than naïve-HFS mice. Fasting insulin was also significantly increased by HFS diet and an NMS/diet interaction with NMS-HFS mice having significantly higher serum insulin levels compared to both NMS-control and naïve-HFS mice (Figure 5C). The homeostatic model assessment of insulin resistance (HOMA-IR) was calculated using fasting glucose and fasting insulin values. HOMA-IR is used to examine insulin resistance where values at or below 1 indicate insulin sensitivity, values between 1.9 and 2.9 indicate early insulin resistance, and values above 2.9 indicate significant insulin resistance. We observed a significant increase in HOMA-IR index due to HFS diet consumption and a significant NMS/diet interaction, resulting in a higher measure in NMS-HFS mice compared to NMS-control mice (Figure 5D). The mean HOMA-IR values for naïve-HFS and NMS-HFS mice were 5.58 ± 1.36 and 9.23 ± 1.32, respectively, indicating significant insulin resistance in these two groups. Taken together, these results suggest that consumption of HFS diet leads to impaired glucose tolerance and insulin resistance. Importantly, the interaction effects of NMS and diet indicate that diet-related alterations in glucose tolerance and insulin resistance are worsened due to NMS exposure, suggesting that NMS mice are more susceptible to diet related changes in glucose tolerance and insulin sensitivity.

**Figure 5.**
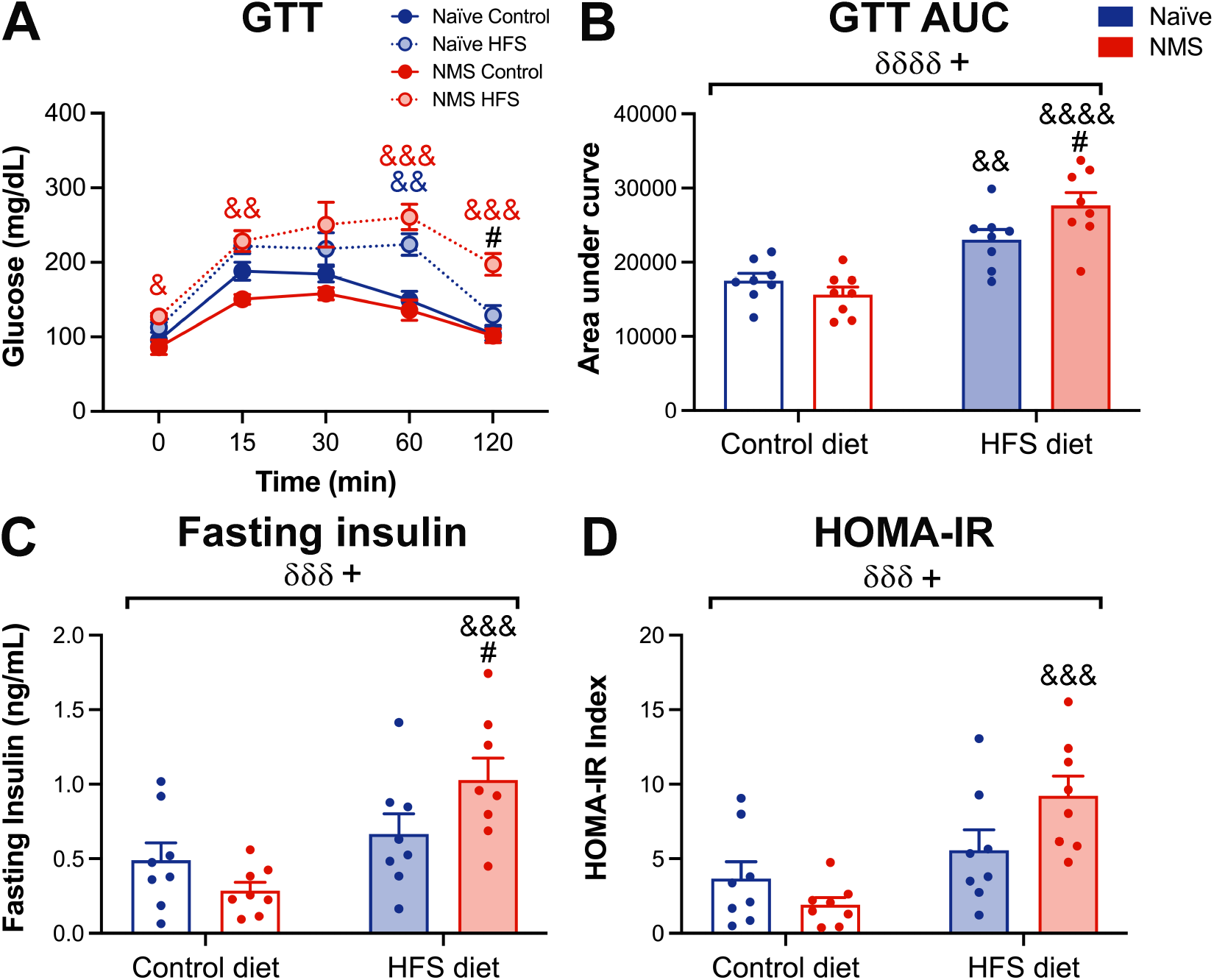
The impact of neonatal maternal separation (NMS) and a high fat/high sucrose (HFS) diet on glucose tolerance, fasting insulin, and HOMA-IR was measured after 25 weeks on the diet. **A**) A significant impact of diet was observed on blood glucose levels over time (*p*<0.0001), as well as a significant interaction effect between NMS and HFS diet (*p*=0.013). NMS-HFS mice had significantly higher blood glucose levels compared to NMS-control at every time point, with the exception of 30-minutes after injection, and were significantly higher than naïve-HFS mice at 120 minutes. Naïve-HFS mice were significantly higher than naïve-control mice only at the 60-minute time point. **B**) Area under the curve (AUC) measurements revealed a significant diet effect (*p*<0.0001) and NMS/diet interaction (*p*=0.0189) with both naïve-HFS and NMS-HFS mice having significantly higher serum glucose compared to their control counterparts. NMS-HFS mice were also significantly higher than naïve-HFS mice. **C**) Fasting insulin levels were also significantly increased due to diet (*p*=0.0006) with an NMS/diet interaction (*p*=0.0248), such that NMS-HFS mice had significantly higher fasting insulin levels compared to both NMS-control and naïve-HFS mice. **D**) HOMA-IR was significantly increased by HFS diet (*p*=0.0003) with an NMS/diet interaction (*p*=0.0237). NMS-HFS mice had significantly higher HOMA-IR compared to NMS-control mice. δ and + denote significant effects of diet and a NMS/diet interaction, respectively, three-way RM ANOVA (**A**), two-way ANOVA (**B-D**). * *p*<0.05 NMS-control vs. naïve-control, &&, &&&, &&&& *p*<0.01, 0.001, 0.0001 HFS vs. control, # *p*<0.05 NMS-HFS vs. naïve-HFS, Fisher’s LSD posttest. n=8 per group.

### Diet and stress increase hepatic fat accumulation, steatosis, and influence glucocorticoid receptor expression

To determine whether NMS and HFS diet resulted in outcomes indicative of non-alcoholic fatty liver disease, liver was examined histologically and for triglyceride levels. Representative images of the liver show increased fat deposition in animals fed a HFS diet, particularly in NMS-HFS mice (Figure 6A). Quantification of liver triglyceride showed a significant impact of diet with NMS-HFS mice having significantly higher levels compared to NMS-control mice (Figure 6B). Although steatosis score was not significantly impacted by NMS or HFS diet, hepatocyte ballooning was significantly increased by NMS (Table 4). Total non-alcoholic fatty liver disease activity (NAS) scores also showed a non-significant increase due to NMS (*p*=0.0791, Table 4).

**Figure 6.**
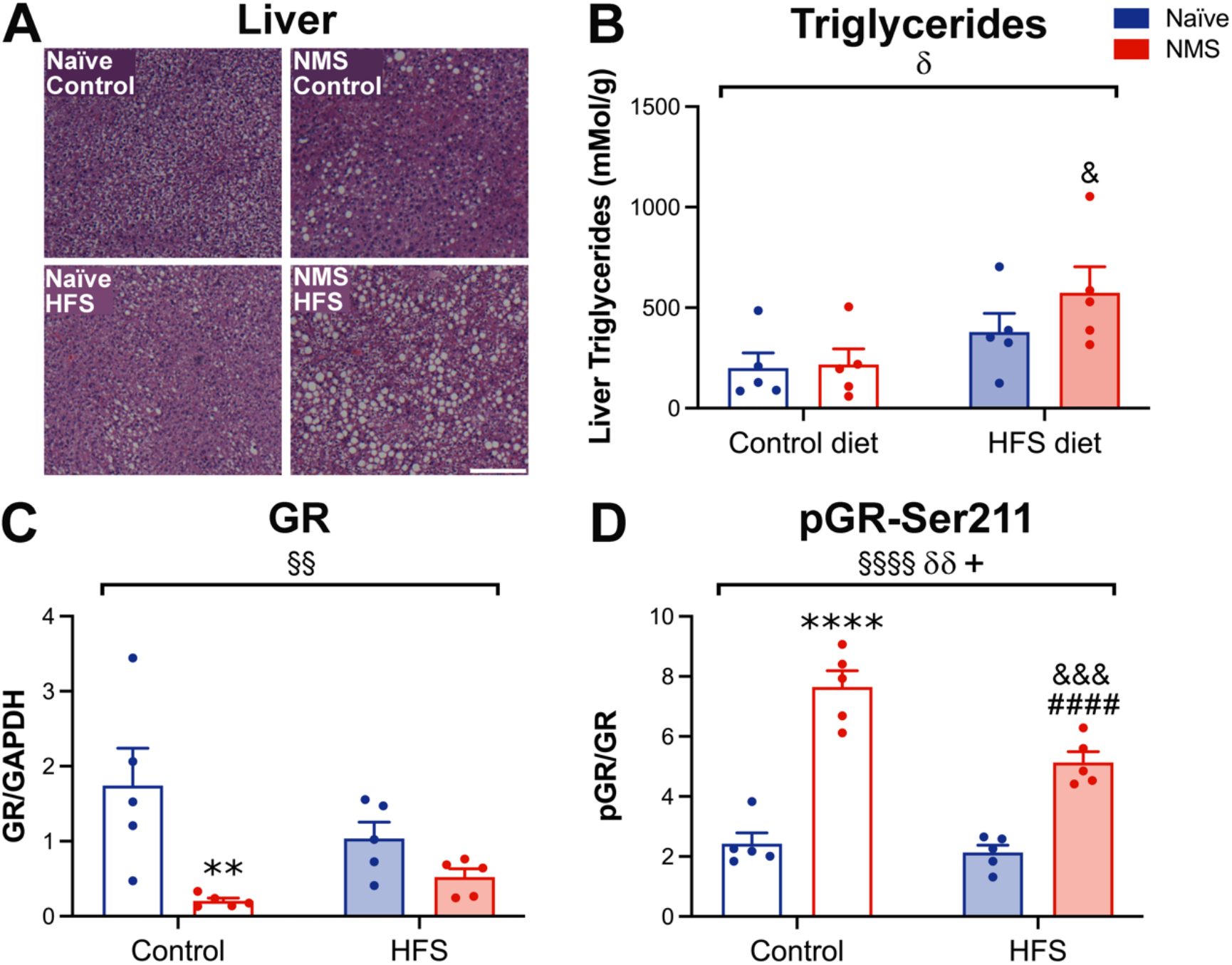
The effect of neonatal maternal separation (NMS) and a high fat/high sucrose (HFS) diet on liver adiposity, triglycerides, and glucocorticoid receptor (GR) protein expression and phosphorylation at Serine residue 211 (pGR-Ser211) was assessed at the end of the study. **A**) Representative hematoxylin and eosin stained liver sections are shown from naïve and NMS mice fed a control or HFS diet. Lipid droplets were noticeably more apparent in liver from NMS-HFS mice. **B**) Measurements of triglyceride content in liver samples revealed a significant increase due to HFS diet (*p*=0.0131), with NMS-HFS mice having a significantly higher level of liver triglyceride compared to NMS-control mice. Western blot of the liver revealed a significant decrease in GR protein levels due to NMS (*p*=0.0019, **C**) and pGR-Ser211 levels were significantly increased by NMS (*p*<0.0001, **D**), decreased by HFS diet (*p*=0.0025), with a significant interaction effect between NMS and HFS (*p*=0.0121). Blots are shown in supplemental figure 1. §, δ, and + denote significant effects of NMS, diet, and a NMS/diet interaction respectively, two-way ANOVA. **, **** *p*<0.01, 0.0001 NMS-control vs. naïve-control, &, &&& *p*<0.05, 0.001 HFS vs. control, Fisher’s LSD posttest. n=5 per group. Scale bar in **A** equals 200μm.

**Table 4.** The impact of neonatal maternal separation (NMS) and high fat/high sucrose (HFS) diet on hepatic steatosis score, hepatocyte ballooning, inflammation, and total non-alcoholic fatty liver disease activity (NAS) score was assessed in H&E-stained liver sections. Steatosis score. Hepatocyte ballooning was significantly increased due to NMS (*p*=0.0195). Total NAS grade was significantly increased by NMS (*p*=0.0018) and HFS diet (*p*=0.0018). § and δ denote significant effects of NMS and diet, respectively, and are indicated by bold text, two-way ANOVA. ** *p*<0.01 NMS-control vs. naïve-control, && *p*<0.01 HFS vs. control, Fisher’s LSD posttest. n=8 per group.

|  | Control Diet |  | HFS Diet |  |
| --- | --- | --- | --- | --- |
|  | Naïve | NMS | Naïve | NMS |
| Steatosis Score | 0.25 ± 0.16 | 0.38 ± 0.18 | 0.38 ± 0.18 | 0.88 ± 0.23 |
| **Hepatocyte Ballooning<sup>§</sup>** | 0 | 0.25 ± 0.16 | 0 | 0.25 ± 0.16 |
| Inflammation | 0 | 0.25 ± 0.25 | 0.12 ± 0.12 | 0.12 ± 0.12 |
| **Total NAS Grade<sup>§</sup>** | 0.25 ± 0.16 | 0.88 ± 0.52 | 0.50 ± 0.27 | 1.25 ± 0.45 |

Stress hormones are known to contribute to the development of steatosis (51). Importantly, phosphorylation of serine 211 on GR has been shown to lead to glucocorticoid hypersensitivity and increased fat deposition in the liver (51). To determine if this mechanism contributes to the steatosis observed in our study, western blot analyses were performed to examine GR levels and the ratio of phosphorylated GR (pGR) to total GR in the liver (Figure 6C and D, Blot shown in supplemental Figure 1). A significant overall effect of NMS was observed on GR protein levels, where NMS exposure led to significantly lower GR expression in the liver (Figure 6C). NMS was also observed to have a significant overall effect on pGR/GR ratio, where NMS exposure led to significantly higher pGR to GR ratio (Figure 6D). pGR/GR ratio was also significantly influenced by diet, with HFS diet consumption resulting in significantly lower pGR/GR ratio. Importantly, a significant NMS x diet interaction was also observed and NMS mice on a control diet were shown to have the highest pGR/GR ratio (Figure 6D). Taken together, these results suggest that NMS and HFS diet synergistically increase fat deposition and measures of steatosis in the liver, possibly through altered glucocorticoid signaling.

### Diet influences fat mass and adipocyte area

To examine how NMS and HFS diet impacted fat distribution, the periovarian and retroperitoneal fat pads were dissected and weighed at time of euthanasia. HFS diet significantly increased both periovarian (Figure *7A*) and retroperitoneal (Figure 7B) fat pad weights. In addition, NMS-HFS mice had significantly heavier periovarian fat pads compared to naïve-HFS mice (Figure 7A). To further characterize differences in periovarian adipose, adipocyte area was measured from H&E stained sections (Figure 7C). Quantification revealed a significant overall effect of diet on adipocyte area with both naïve-HFS and NMS-HFS mice having significantly larger periovarian adipocytes compared to control diet-fed mice (Figure 7D).

**Figure 7.**
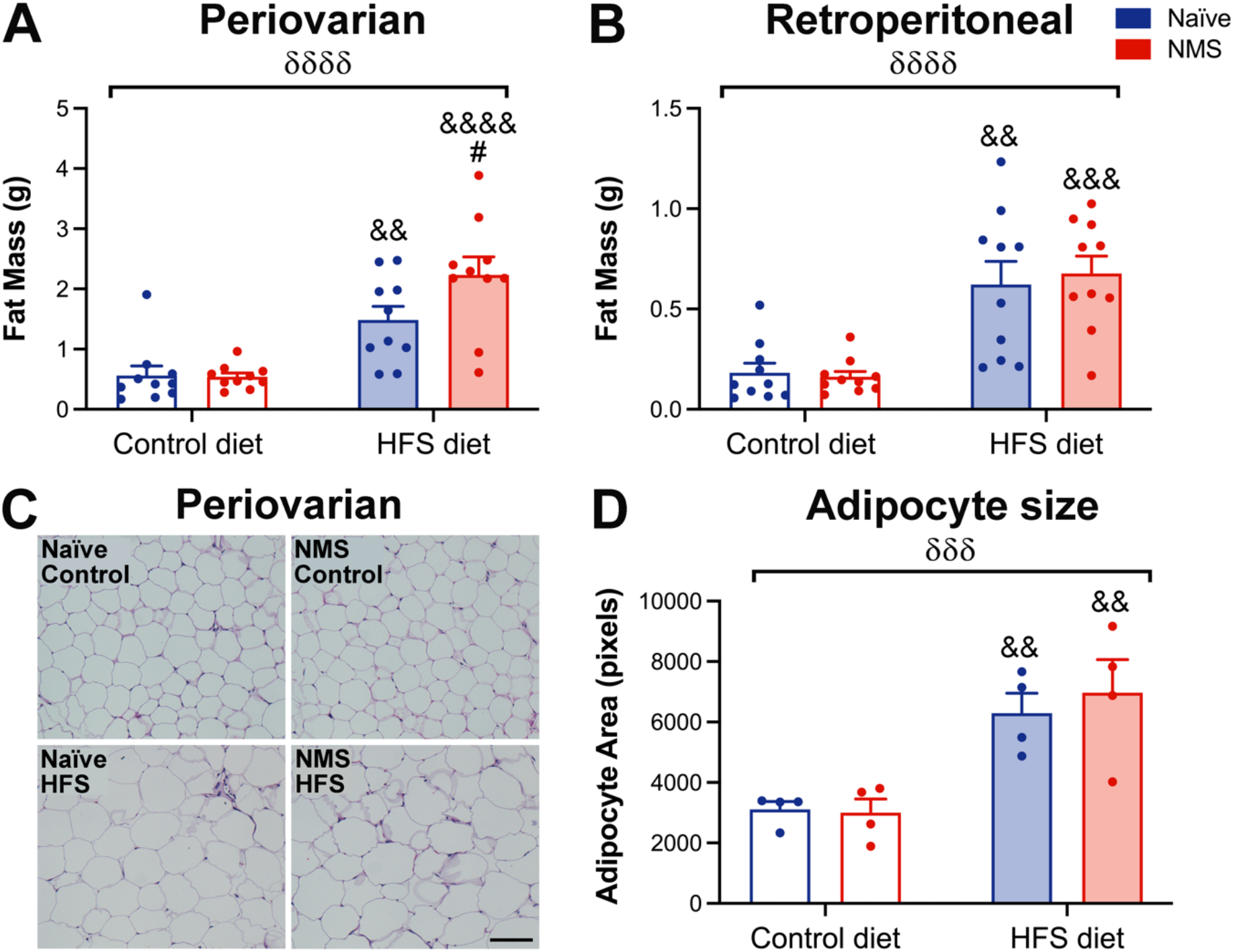
The impact of neonatal maternal separation (NMS) and a high fat/high sucrose (HFS) diet on periovarian and retroperitoneal adipose was determined at the end of the study. **A)** Periovarian fat weight was significantly increased by HFS diet (*p*<0.0001) with naïve-HFS and NMS-HFS mice having significantly heavier fat pads compared to control diet-fed counterparts. NMS-HFS periovarian fat was also significantly heavier than that from naïve-HFS mice. **B)** Retroperitoneal fat weight was also significantly increased by HFS diet (*p*<0.0001) with naïve-HFS and NMS-HFS mice having significantly heavier fat pads compared to control diet-fed counterparts. **C)** Periovarian fat pads were stained with hematoxylin and eosin and adipocyte size was measured. Periovarian adipocyte area was significantly increased by HFS diet (*p*=0.0002) with naïve-HFS and NMS-HFS mice having significantly larger adipocytes compared to control diet-fed counterparts. δ denotes a significant effect of diet, two-way ANOVA. &&, &&&, &&&& *p*<0.01, 0.001, 0.0001 HFS vs. control, # *p*<0.05 NMS-HFS vs. naïve-HFS, Fisher’s LSD posttest. n=10 (**A-B**) and n=4 (**C-D**) per group. Scale bar in **C** equals 100μm.

### Diet alters periovarian adipose gene expression

Periovarian adipose tissue was further examined for changes in gene expression that may be contributing to/resulting from the metabolic and behavioral differences observed between our groups. Several macrophage markers were examined due to their association to chronic pain syndromes (52) and high fat diet consumption (53). A significant overall effect of diet was observed on the expression of the macrophage markers F4/80, driven specifically by increases in NMS-HFS mice, and CD68, which was increased in both naïve-HFS and NMS-HFS mice, compared to control (Figure 8A). To examine macrophage inflammatory status, expression of CD11b and CD11c were analyzed. Diet significantly increased both CD11b and CD11c expression, with specific increases in naïve-HFS mice (Figure 8A). No overall effects were observed on TNFa, IL10, or tryptase, although naïve-HFS mice had significantly higher TNFa mRNA levels compared to naïve-control mice (Figure 8B).

**Figure 8.**
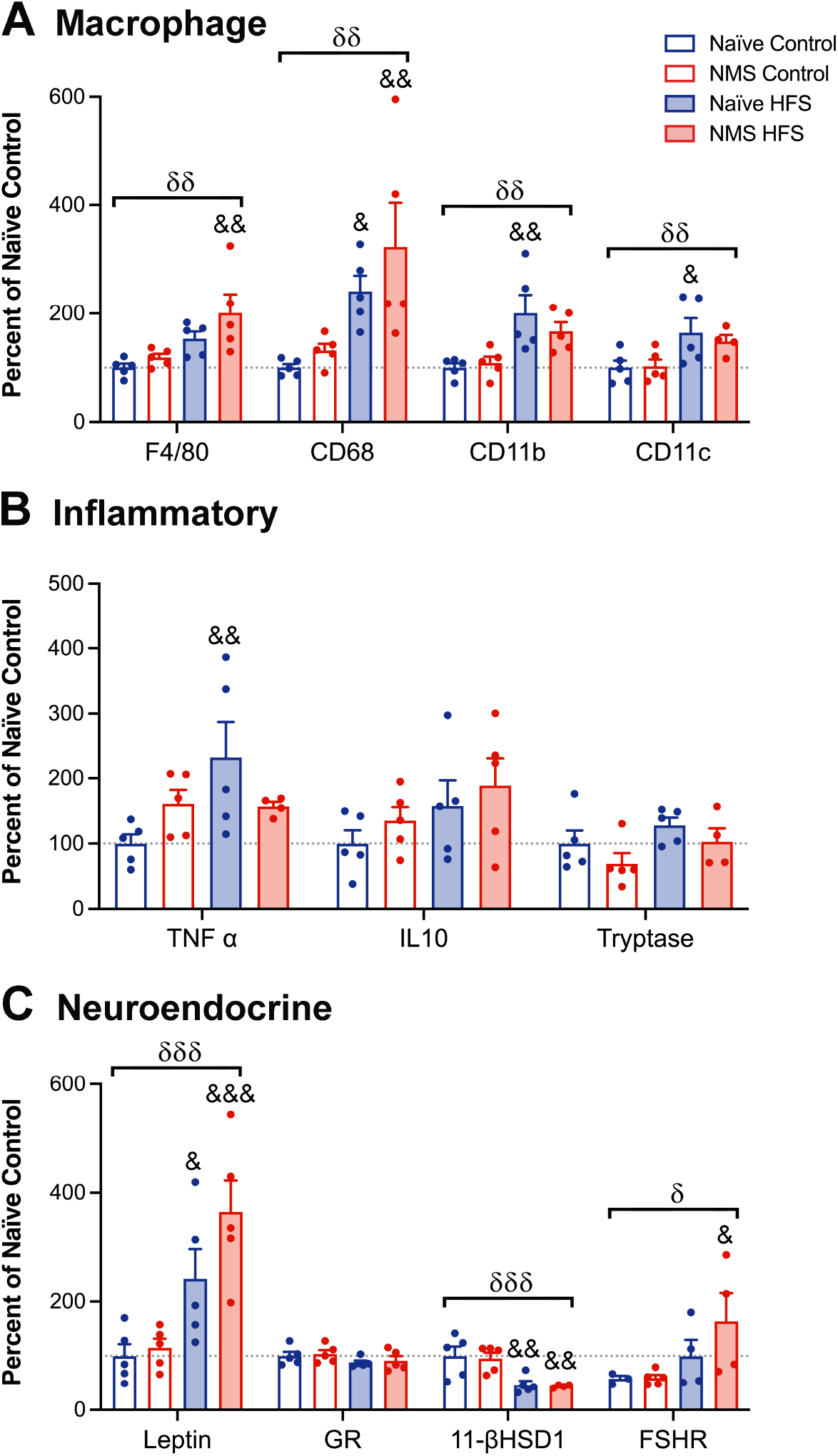
The effect of neonatal maternal separation (NMS) and high fat/high sucrose (HFS) diet on gene expression of inflammatory and neuroendocrine markers in periovarian adipose tissue was measured at the end of the study. **A**) A significant effect of HFS diet on increasing F4/80 (*p*=0.0026), CD68 (*p*=0.0015), CD11b (*p*=0.0011), and CD11c (*p*=0.0082) mRNA levels was observed. NMS-HFS mice had increased F4/80 and CD68, whereas naïve-HFS mice had increased CD68, CD11b, and CD11c, compared to their control diet-fed counterparts. **B**) TNFa mRNA levels were significantly increased in naïve-HFS mice compared to naïve-control. No significant effects of NMS or HFS diet were observed on IL10 or tryptase mRNA levels. **C**) HFS diet significantly increased mRNA levels of leptin (*p*=0.0011) and FSHR (*p*=0.0351), whereas it decreased levels of 11-βHSD1 (*p*=0.0004). δ denotes a significant effect of diet, two-way ANOVA. &, &&, &&& *p*<0.05, 0.01, 0.001, HFS vs. control, Fisher’s LSD posttest. n=5 per group.

Perigonadal adipose tissue acts as an important endocrine organ that both secretes and is responsive to hormonal signals (54, 55). To analyze possible changes in neuroendocrine signaling, the gene expression of leptin, 11β-hydroxysteroid dehydrogenase type 1 (11βHSD1), GR, and follicle-stimulating hormone receptor (FSHR) were measured. A significant overall effect of diet was observed on leptin expression, such that both HFS diet-fed groups had significantly higher leptin mRNA levels compared to control diet-fed groups (Figure 8C). Levels of 11βHSD1, which converts inactive glucocorticoids to their active form (56), were significantly reduced in HFS diet-fed groups, whereas FSHR was significantly increased by HFS diet, particularly in NMS-HFS mice (Figure 8C). Neither diet nor NMS had a significant effect of GR expression (Figure 8C). In addition, we observed a trend toward increased serum corticosterone due to NMS (*p*=0.0703) and a non-significant decrease in serum protein levels of FSH in NMS-control mice compared to naïve-control mice (*p*=0.0647, Supplemental Figure 2).

## Discussion

The development of obesity is highly complex, with genetic, physiologic, environmental, and life-style factors all contributing to the condition (57–60). The consumption of a HFS diet can lead to weight gain, increased adiposity, and impaired glucose tolerance (10, 61–64), but previous results show that female mice are generally protected from these effects compared to males (65, 66). Similarly, exposure to stressful events early in life is correlated with increased body weight and poor metabolic outcomes (15, 67–70). HFS diet consumption and ELS exposure are both highly prevalent in the U.S. (1, 2, 9, 18) and often co-occur within populations (29, 30, 71). Therefore, it is imperative to understand both the individual and combined effects of ELS and HFS diet consumption on long term metabolic outcomes. Here, we examined the combined effects of NMS and HFS diet consumption on body weight, adiposity, glucose tolerance, steatosis, energy intake, energy expenditure, and pain-like behavior, in female mice.

Consistent with our previous studies (34, 48, 50, 72, 73), we observed mechanical hypersensitivity of the hindpaw and perigenital region due to NMS exposure. These results confirm the NMS phenotype within this study. Additionally, obesity and metabolic disorders often coincide with chronic pain syndromes and have been associated with widespread low-grade inflammation (74, 75). In the current study we also observed a significant effect of HFS diet worsening mechanical hypersensitivity. We previously made a similar observation in NMS mice fed a diet high in omega-3 fatty acids (35%) and no sucrose (50). Supplementation of the diet with anti-inflammatory components improved mechanical sensitivity, although it had no impact on (and in some cases increased) perigonadal macrophage and inflammatory markers. A similar observation of increased macrophage-related genes in perigonadal adipose was made in the current study, suggesting that increased inflammation contributes to the widespread allodynia phenotype. Importantly, we observed an NMS and HFS diet interaction on increasing hindpaw mechanical hypersensitivity, suggesting the co-occurrence of ELS and a Western diet likely increases the risk of chronic pain syndromes.

HFS diet consumption in female mice resulted in increased body weight and percent body fat, although its effects were observed earlier in NMS mice compared to naïve. Our findings showed that, after 10-weeks, female NMS mice had significantly higher body weight and percent body fat compared to naïve mice, across both diets. The distinction between NMS and naïve mice was lost at the 20-week timepoint, although HFS diet continued to increase both outcomes. Differences in feeding behavior likely contributed to the earlier gains in NMS-related body weight and adiposity, as NMS mice had a higher average food consumption during the first 10 weeks on the diet. Food intake and feed efficiency were increased only by HFS diet in the latter half of the study, potentially due to the increased energy requirements that accompany larger body mass. These results are consistent with our previous study (67) and with Bernardi et al. (76), which both showed an ELS-related increase in food consumption accompanied by increased body weight in ELS exposed animals. Overall, these data show that NMS removes protection against early but not long-term weight gain that has been reported in female mice.

Although NMS did not have a statistically significant impact on body weight at the 20-week timepoint, there was a lasting effect on energy expenditure. Specifically, exposure to NMS significantly increased TEE and REE, while diet increased TEE, REE, and EI and decreased NREE. When the energy expenditure data were adjusted for the observed differences in body weight, the effects of both NMS and diet were lost, suggesting that body weight was driving energy intake and expenditure. This outcome is particularly interesting because we did not observe a significant effect of NMS on body weight at the 20-week timepoint. Therefore, although NMS did not lead to a difference in body weight, it did result in a biologically relevant difference as the NMS related changes in body weight were primarily responsible for the NMS effect on TEE and REE. Future work will focus on differences in energy intake and expenditure at an earlier timepoint, coinciding with a more rapid rate of weight gain, to determine their contributions to NMS-related weight gain

Obesity and diabetes are often observed as comorbid disorders associated with HFS diet consumption and ELS (9, 77–80). The ‘developmental origins of adult disease’ hypothesis identified a relationship between adverse experiences in early life and poor health outcomes such as diabetes and coronary heart disease in adulthood (81). These studies emphasize the importance of the early life environment in programing long term health outcomes. We observed a significant interaction effect of NMS and HFS diet resulting in impaired glucose tolerance, elevated fasting insulin, and higher HOMA-IR index values, all of which are consistent with insulin resistance. Importantly, these results indicate that NMS exposure results in increased susceptibility to diet related changes in glucose tolerance. Considering that estrogen has a protective effect on glucose tolerance (82, 83), it is likely that the combined effect of NMS and HFS diet were required to overcome the effects of estrogen. These results are consistent with other studies indicating that ELS exposure increases risk for diabetes in adulthood (15, 84).

Obesity and diabetes have also been determined to increase the risk of developing nonalcoholic fatty liver disease. Of note, our group and others have repeatedly shown that females are protected from hepatic steatosis, a protection that goes away with experimental or natural loss of ovarian function (85, 86). Glucocorticoids have also been documented to stimulate free fatty acid release from adipose tissue resulting in increased ectopic lipid deposition in the liver (87, 88). Importantly, the phosphorylation status of the GR modulates its sensitivity to glucocorticoids ultimately influencing the level of steatosis. Although we only saw diet related differences in liver triglyceride levels, we observed an effect of NMS on increasing NAS score and altering hepatic GR protein content and phosphorylation. NMS mice had lower GR protein levels, but a significantly higher ratio of pGR/GR. These results suggest that the increased sensitivity of pGR may have compensated for the reduced GR protein content in NMS mice, resulting in higher NAS scores. The GR phosphorylation observed in NMS mice may have been driven by differences in follicle stimulating hormone (FSH) levels. Quinn and colleagues determined that elevated FSH levels due to ovariectomy directly results in GR phosphorylation, glucocorticoid hypersensitivity, and increased steatosis (51). Considering that glucocorticoids stimulate FSH secretion (89, 90) and chronic stress exposure has been associated with elevated FSH levels (91, 92); it is possible that NMS-related changes in FSH secretion may contribute to the increased GR phosphorylation. Although we did not observe any significant changes in serum FSH content, HFS diet significantly increased FSHR expression in periovarian adipose, particularly in NMS mice. Future studies will investigate the effects of NMS and HFS diet on FSHR expression and signaling pathways in the liver of female mice to determine potential sex-specific interactions between estrogen, FSH, and GR.

Body composition and distribution of adipose tissue are both important predictors of metabolic health (93, 94) as is ectopic storage of lipids in the liver. Additionally, adipose tissue is known to play an important endocrine role, which can be disrupted due to HFS diet and fat accumulation (54, 55). Adipocytes secrete leptin, which influences the regulation of body fat through the promotion of satiety (95, 96) and contributes to obesity-related inflammation via stimulation of cytokine release (54, 97–99). Chronically elevated leptin levels, due to increased adiposity, can eventually lead to leptin resistance and may be involved in obesity-related hyperphagia and inflammation (100, 101). Here, we observed a significant elevation in leptin expression in adipose tissue due to high fat feeding. This difference in leptin expression may have contributed to the inflammation and increased caloric intake observed in our study. Although we did not observe an effect of NMS on leptin expression in adulthood, it is possible that leptin levels may have been altered early in life, potentially contributing to alterations in hypothalamic development and NMS-related hyperphagia. Several studies have indicated that ELS exposure results in reduced leptin levels early in life and ultimately may alter hypothalamic development (102, 103). Early life changes in hypothalamic development have been documented to persist into adulthood and to have long-term effects on energy regulation, potentially contributing to adult obesity (104–111). Future work will further examine hypothalamic development and appetite regulation after NMS exposure.

ELS exposure has been documented to lead to other hypothalamic changes including dysregulation of the hypothalamic-pituitary-adrenal axis resulting in increased glucocorticoid secretion (72, 112). Similarly, obesity has been linked to high glucocorticoid levels (87, 113, 114), which can increase adipogenesis and visceral fat deposition (87, 88). Therefore, differences in glucocorticoid secretion and responsiveness may contribute to ELS-related weight gain. Obese patients have been documented to have elevated GR and 11-βHSD1, which converts glucocorticoids to their active form (115, 116). Although we did not see any significant differences in GR expression, we did observe a significant reduction in 11-βHSD1 expression due to HFS diet consumption. These results are consistent with other studies which also observed down regulation of 11 -βHSD1 due to high fat feeding (67, 117, 118).

Taken together our results indicate that NMS and HFS diet consumption both contribute to metabolic dysfunction and mechanical hypersensitivity in female mice. The NMS and HFS diet effects and interactions suggest that female mice exposed to NMS are more susceptible to diet-induced weight gain, adiposity, hepatic steatosis, and glucose intolerance which are associated with glucocorticoid signaling capacity. NMS-related hyperphagia likely contributes to these outcomes although additional work is needed to identify the underlying mechanisms driving this condition.

## Acknowledgments

We would like to thank Drs. Kyle Baumbauer, Paige Geiger, and Doug Wright for their thoughtful contributions towards the development, execution, and interpretation of this project.

---

**Supplemental Figure 1.**
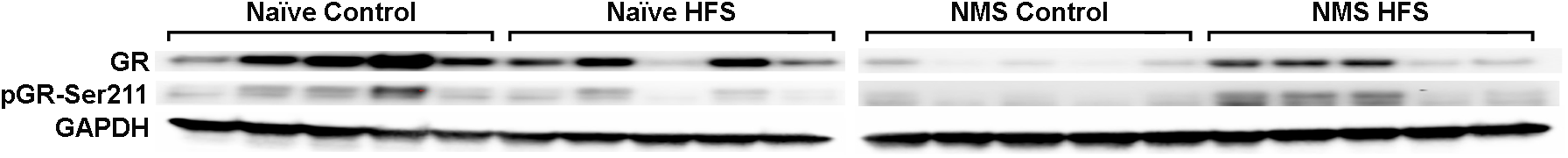
Western blots for data shown in Figure 7. n=5 per group.

**Supplemental Figure 2.**
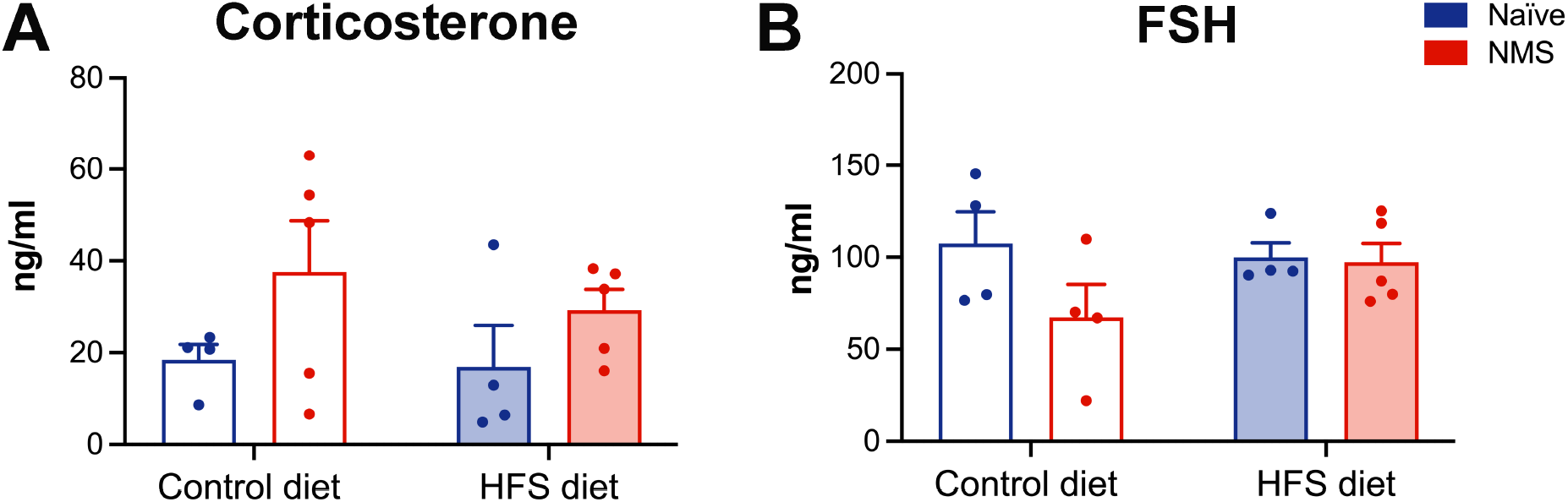
The effect of neonatal maternal separation (NMS) and high fat/high sucrose (HFS) diet on serum protein levels of corticosterone and follicle stimulating hormone (FSH) were measured at the end of the study. **A**) A trend toward increased corticosterone was observed for NMS (*p*=0.0703) with no impact of diet. **B**) No overall effects of NMS or HFS diet were observed on FSH, however a non-significant decrease in NMS-control compared to naïve-control (*p*=0.0647) was measured. Two-way ANOVA, Fisher’s LSD posttest. n=4-5 per group.

## References

1. Hales CM, Carroll MD, Fryar CD, and Ogden CL. Prevalence of Obesity and Severe Obesity Among Adults: United States, 2017-2018. NCHS Data Brief 1–8, 2020.

2. Hales CM CM, Fryar CD, Ogden CL. Prevalence of obesity among adults and youth: United States, 2015-2016. NCHS data brief, no288. Hyattsville, MD: National Center for Health Statistics. 2017.

3. Kannel WB, Cupples LA, Ramaswami R, Stokes J, Kreger BE, and Higgins M. Regional obesity and risk of cardiovascular disease; the Framingham Study. J Clin Epidemiol 44: 183–190, 1991.

4. Lakka HM, Lakka TA, Tuomilehto J, and Salonen JT. Abdominal obesity is associated with increased risk of acute coronary events in men. Eur Heart J 23: 706–713, 2002.

5. Lakka HM, Salonen JT, Tuomilehto J, Kaplan GA, and Lakka TA. Obesity and weight gain are associated with increased incidence of hyperinsulinemia in non-diabetic men. Hormone and metabolic research = Hormon-und Stoffwechselforschung = Hormones et metabolisme 34: 492–498, 2002.

6. Lambert AA, Putcha N, Drummond MB, Boriek AM, Hanania NA, Kim V, Kinney GL, McDonald MN, Brigham EP, Wise RA, McCormack MC, Hansel NN, and Investigators CO. Obesity Is Associated With Increased Morbidity in Moderate to Severe COPD. Chest 151: 68–77, 2017.

7. Steele CB, Thomas CC, Henley SJ, Massetti GM, Galuska DA, Agurs-Collins T, Puckett M, and Richardson LC. Vital Signs: Trends in Incidence of Cancers Associated with Overweight and Obesity - United States, 2005-2014. MMWR Morb Mortal Wkly Rep 66: 1052–1058, 2017.

8. Kim DD, and Basu A. Estimating the Medical Care Costs of Obesity in the United States: Systematic Review, Meta-Analysis, and Empirical Analysis. Value Health 19: 602–613, 2016.

9. Felitti VJ, Anda RF, Nordenberg D, Williamson DF, Spitz AM, Edwards V, Koss MP, and Marks JS. Relationship of childhood abuse and household dysfunction to many of the leading causes of death in adults. The Adverse Childhood Experiences (ACE) Study. American journal of preventive medicine 14: 245–258, 1998.

10. Middlebrooks JS. The effects of childhood stress on health across a lifespan. edited by Audage NC. Atlanta (GA): Centers for Disease Control and Prevention, National Center for Injury Prevention and Control: 2008.

11. Davis DA, Luecken LJ, and Zautra AJ. Are reports of childhood abuse related to the experience of chronic pain in adulthood? A meta-analytic review of the literature. The Clinical journal of pain 21: 398–405, 2005.

12. Alvarez J, Pavao J, Baumrind N, and Kimerling R. The relationship between child abuse and adult obesity among california women. American journal of preventive medicine 33: 28–33, 2007.

13. Miller AL, and Lumeng JC. Pathways of Association from Stress to Obesity in Early Childhood. Obesity 26: 1117–1124, 2018.

14. Richardson AS, Dietz WH, and Gordon-Larsen P. The association between childhood sexual and physical abuse with incident adult severe obesity across 13 years of the National Longitudinal Study of Adolescent Health. Pediatr Obes 9: 351–361, 2014.

15. Williamson DF, Thompson TJ, Anda RF, Dietz WH, and Felitti V. Body weight and obesity in adults and self-reported abuse in childhood. Int J Obes Relat Metab Disord 26: 1075–1082, 2002.

16. Noll JG, Zeller MH, Trickett PK, and Putnam FW. Obesity risk for female victims of childhood sexual abuse: a prospective study. Pediatrics 120: e61–67, 2007.

17. Midei AJ, and Matthews KA. Interpersonal violence in childhood as a risk factor for obesity: a systematic review of the literature and proposed pathways. Obes Rev 12: e159–172, 2011.

18. U.S. Department of Health and Human Services AfCaF AoC, Youth and Families, Children’s Bureau. Child maltreatment. 2018.

19. Knickmeyer RC, Gouttard S, Kang C, Evans D, Wilber K, Smith JK, Hamer RM, Lin W, Gerig G, and Gilmore JH. A structural MRI study of human brain development from birth to 2 years. J Neurosci 28: 12176–12182, 2008.

20. Mainardi M, Scabia G, Vottari T, Santini F, Pinchera A, Maffei L, Pizzorusso T, and Maffei M. A sensitive period for environmental regulation of eating behavior and leptin sensitivity. Proceedings of the National Academy of Sciences of the United States of America 107: 16673–16678, 2010.

21. Bouret SG, and Simerly RB. Minireview: Leptin and development of hypothalamic feeding circuits. Endocrinology 145: 2621–2626, 2004.

22. Bouret SG. Role of early hormonal and nutritional experiences in shaping feeding behavior and hypothalamic development. J Nutr 140: 653–657, 2010.

23. Remmers F, and Delemarre-van de Waal HA. Developmental programming of energy balance and its hypothalamic regulation. Endocrine reviews 32: 272–311, 2011.

24. Medina-Remon A, Kirwan R, Lamuela-Raventos RM, and Estruch R. Dietary patterns and the risk of obesity, type 2 diabetes mellitus, cardiovascular diseases, asthma, and neurodegenerative diseases. Crit Rev Food Sci Nutr 58: 262–296, 2018.

25. Tremblay A, Plourde G, Despres JP, and Bouchard C. Impact of dietary fat content and fat oxidation on energy intake in humans. Am J Clin Nutr 49: 799–805, 1989.

26. Romieu I, Willett WC, Stampfer MJ, Colditz GA, Sampson L, Rosner B, Hennekens CH, and Speizer FE. Energy intake and other determinants of relative weight. Am J Clin Nutr 47: 406–412, 1988.

27. Committee DGA. Report of the Dietary Guidelines Advisory Committee on the Dietary Guidelines for Americans, 2010 Washington, DC: U.S.: Department of Health and Human Services, U.S. Department of Agriculture, 2010.

28. Darmon N, Ferguson E, and Briend A. Do economic constraints encourage the selection of energy dense diets? Appetite 41: 315–322, 2003.

29. Lehman BJ, Taylor SE, Kiefe CI, and Seeman TE. Relation of childhood socioeconomic status and family environment to adult metabolic functioning in the CARDIA study. Psychosomatic medicine 67: 846–854, 2005.

30. Taylor SE, Lerner JS, Sage RM, Lehman BJ, and Seeman TE. Early environment, emotions, responses to stress, and health. J Pers 72: 1365–1393, 2004.

31. Lovejoy JC, Sainsbury A, and Stock Conference Working G. Sex differences in obesity and the regulation of energy homeostasis. Obes Rev 10: 154–167, 2009.

32. Hong J, Stubbins RE, Smith RR, Harvey AE, and Nunez NP. Differential susceptibility to obesity between male, female and ovariectomized female mice. Nutr J 8: 11, 2009.

33. Chen JL, Guo J, Mao P, Yang J, Jiang S, He W, Lin CX, and Lien K. Are the factors associated with overweight/general obesity and abdominal obesity different depending on menopausal status? PloS one 16: e0245150, 2021.

34. Fuentes IM, Pierce AN, O’Neil PT, and Christianson JA. Assessment of Perigenital Sensitivity and Prostatic Mast Cell Activation in a Mouse Model of Neonatal Maternal Separation. Journal of visualized experiments: JoVE e53181, 2015.

35. Dixon WJ. Efficient analysis of experimental observations. Annual review of pharmacology and toxicology 20: 441–462, 1980.

36. Chaplan SR, Bach FW, Pogrel JW, Chung JM, and Yaksh TL. Quantitative assessment of tactile allodynia in the rat paw. Journal of neuroscience methods 53: 55–63, 1994.

37. Tai MM. A mathematical model for the determination of total area under glucose tolerance and other metabolic curves. Diabetes Care 17: 152–154, 1994.

38. Parlee SD, Lentz SI, Mori H, and MacDougald OA. Quantifying size and number of adipocytes in adipose tissue. Methods in enzymology 537: 93–122, 2014.

39. Kleiner DE, Brunt EM, Van Natta M, Behling C, Contos MJ, Cummings OW, Ferrell LD, Liu YC, Torbenson MS, Unalp-Arida A, Yeh M, McCullough AJ, Sanyal AJ, and Network NSCR. Design and validation of a histological scoring system for nonalcoholic fatty liver disease. Hepatology 41: 1313–1321, 2005.

40. Ramakers C, Ruijter JM, Deprez RH, and Moorman AF. Assumption-free analysis of quantitative real-time polymerase chain reaction (PCR) data. Neuroscience letters 339: 62–66, 2003.

41. Pfaffl MW. A new mathematical model for relative quantification in real-time RT-PCR. Nucleic acids research 29: e45, 2001.

42. Tramullas M, Finger BC, Dinan TG, and Cryan JF. Obesity Takes Its Toll on Visceral Pain: High-Fat Diet Induces Toll-Like Receptor 4-Dependent Visceral Hypersensitivity. PloS one 11: e0155367, 2016.

43. Brandao AF, Bonet IJM, Pagliusi M, Jr., Zanetti GG, Pho N, Tambeli CH, Parada CA, Vieira AS, and Sartori CR. Physical Activity Induces Nucleus Accumbens Genes Expression Changes Preventing Chronic Pain Susceptibility Promoted by High-Fat Diet and Sedentary Behavior in Mice. Frontiers in neuroscience 13: 1453, 2019.

44. Green PG, Chen X, Alvarez P, Ferrari LF, and Levine JD. Early-life stress produces muscle hyperalgesia and nociceptor sensitization in the adult rat. Pain 152: 2549–2556, 2011.

45. Sachs-Ericsson NJ, Sheffler JL, Stanley IH, Piazza JR, and Preacher KJ. When Emotional Pain Becomes Physical: Adverse Childhood Experiences, Pain, and the Role of Mood and Anxiety Disorders. J Clin Psychol 73: 1403–1428, 2017.

46. Brennenstuhl S, and Fuller-Thomson E. The Painful Legacy of Childhood Violence: Migraine Headaches Among Adult Survivors of Adverse Childhood Experiences. Headache 55: 973–983, 2015.

47. Pierce AN, Di Silvestro ER, Eller OC, Wang R, Ryals JM, and Christianson JA. Urinary bladder hypersensitivity and dysfunction in female mice following early life and adult stress. Brain Res 1639: 58–73, 2016.

48. Fuentes IM, Pierce AN, Di Silvestro ER, Maloney MO, and Christianson JA. Differential Influence of Early Life and Adult Stress on Urogenital Sensitivity and Function in Male Mice. Front Syst Neurosci 11: 97, 2017.

49. Eller OC, Yang X, Fuentes IM, Pierce AN, Jones BM, Brake AD, Wang R, Dussor G, and Christianson JA. Voluntary Wheel Running Partially Attenuates Early Life Stress-Induced Neuroimmune Measures in the Dura and Evoked Migraine-Like Behaviors in Female Mice. Front Physiol 12: 665732, 2021.

50. Eller OC, Foright RM, Brake AD, Winter MK, Bantis LE, Morris EM, Thyfault JP, and Christianson JA. An Omega-3-rich Anti-inflammatory Diet Improved Widespread Allodynia and Worsened Metabolic Outcomes in Adult Mice Exposed to Neonatal Maternal Separation. Neuroscience 468: 53–67, 2021.

51. Quinn MA, Xu X, Ronfani M, and Cidlowski JA. Estrogen Deficiency Promotes Hepatic Steatosis via a Glucocorticoid Receptor-Dependent Mechanism in Mice. Cell reports 22: 2690–2701, 2018.

52. Ristoiu V. Contribution of macrophages to peripheral neuropathic pain pathogenesis. Life sciences 93: 870–881, 2013.

53. Fink LN, Costford SR, Lee YS, Jensen TE, Bilan PJ, Oberbach A, Bluher M, Olefsky JM, Sams A, and Klip A. Pro-inflammatory macrophages increase in skeletal muscle of high fat-fed mice and correlate with metabolic risk markers in humans. Obesity 22: 747–757, 2014.

54. Fontana L, Eagon JC, Trujillo ME, Scherer PE, and Klein S. Visceral fat adipokine secretion is associated with systemic inflammation in obese humans. Diabetes 56: 1010–1013, 2007.

55. Scherer PE. The Multifaceted Roles of Adipose Tissue-Therapeutic Targets for Diabetes and Beyond: The 2015 Banting Lecture. Diabetes 65: 1452–1461, 2016.

56. Tomlinson JW, Walker EA, Bujalska IJ, Draper N, Lavery GG, Cooper MS, Hewison M, and Stewart PM. 11beta-hydroxysteroid dehydrogenase type 1: a tissue-specific regulator of glucocorticoid response. Endocrine reviews 25: 831–866, 2004.

57. Hill JO, Wyatt HR, Reed GW, and Peters JC. Obesity and the environment: where do we go from here? Science 299: 853–855, 2003.

58. Wright SM, and Aronne LJ. Causes of obesity. Abdom Imaging 37: 730–732, 2012.

59. Herrera BM, Keildson S, and Lindgren CM. Genetics and epigenetics of obesity. Maturitas 69: 41–49, 2011.

60. Schrauwen P, and Westerterp KR. The role of high-fat diets and physical activity in the regulation of body weight. Br J Nutr 84: 417–427, 2000.

61. Hariri N, and Thibault L. High-fat diet-induced obesity in animal models. Nutr Res Rev 23: 270–299, 2010.

62. Collins S, Martin TL, Surwit RS, and Robidoux J. Genetic vulnerability to diet-induced obesity in the C57BL/6J mouse: physiological and molecular characteristics. Physiology & behavior 81: 243–248, 2004.

63. Surwit RS, Kuhn CM, Cochrane C, McCubbin JA, and Feinglos MN. Diet-induced type II diabetes in C57BL/6J mice. Diabetes 37: 1163–1167, 1988.

64. Swinburn BA, Caterson I, Seidell JC, and James WP. Diet, nutrition and the prevention of excess weight gain and obesity. Public Health Nutr 7: 123–146, 2004.

65. Lovejoy JC, Sainsbury A, and Group SCW. Sex differences in obesity and the regulation of energy homeostasis. Obes Rev 10: 154–167, 2009.

66. Hong J, Stubbins RE, Smith RR, Harvey AE, and Núñez NP. Differential susceptibility to obesity between male, female and ovariectomized female mice. Nutr J 8: 11, 2009.

67. Eller OC, Morris EM, Thyfault JP, and Christianson JA. Early life stress reduces voluntary exercise and its prevention of diet-induced obesity and metabolic dysfunction in mice. Physiology & behavior 223: 113000, 2020.

68. Delpierre C, Fantin R, Barboza-Solis C, Lepage B, Darnaudery M, and Kelly-Irving M. The early life nutritional environment and early life stress as potential pathways towards the metabolic syndrome in mid-life? A lifecourse analysis using the 1958 British Birth cohort. BMC public health 16: 815, 2016.

69. Hales CN, and Barker DJ. Type 2 (non-insulin-dependent) diabetes mellitus: the thrifty phenotype hypothesis. Diabetologia 35: 595–601, 1992.

70. Fall CH, Osmond C, Barker DJ, Clark PM, Hales CN, Stirling Y, and Meade TW. Fetal and infant growth and cardiovascular risk factors in women. BMJ (Clinical research ed 310: 428–432, 1995.

71. Finkelhor D, Shattuck A, Turner H, and Hamby S. A revised inventory of Adverse Childhood Experiences. Child abuse & neglect 48: 13–21, 2015.

72. Pierce AN, Ryals JM, Wang R, and Christianson JA. Vaginal hypersensitivity and hypothalamic-pituitary-adrenal axis dysfunction as a result of neonatal maternal separation in female mice. Neuroscience 263: 216–230, 2014.

73. Fuentes IM, Jones BM, Brake AD, Pierce AN, Eller OC, Supple RM, Wright DE, and Christianson JA. Voluntary wheel running improves outcomes in an early life stress-induced model of urologic chronic pelvic pain syndrome in male mice. Pain 162: 1681–1691, 2021.

74. Fink LN, Costford SR, Lee YS, Jensen TE, Bilan PJ, Oberbach A, Blüher M, Olefsky JM, Sams A, and Klip A. Pro-inflammatory macrophages increase in skeletal muscle of high fat-fed mice and correlate with metabolic risk markers in humans. Obesity 22: 747–757, 2014.

75. Okifuji A, and Hare BD. The association between chronic pain and obesity. J Pain Res 8: 399–408, 2015.

76. Bernardi JR, Ferreira CF, Senter G, Krolow R, de Aguiar BW, Portella AK, Kauer-Sant’anna M, Kapczinski F, Dalmaz C, Goldani MZ, and Silveira PP. Early life stress interacts with the diet deficiency of omega-3 fatty acids during the life course increasing the metabolic vulnerability in adult rats. PloS one 8: e62031, 2013.

77. Astrup AV, Madsbad S, and Finer N. [New discoveries about the cause of diabetes. Type 2 diabetes mellitus changed to “obesity-dependent diabetes mellitus”]. Ugeskr Laeger 163: 141–143, 2001.

78. Leong KS, and Wilding JP. Obesity and diabetes. Baillieres Best Pract Res Clin Endocrinol Metab 13: 221–237, 1999.

79. Felitti VJ. [The relationship of adverse childhood experiences to adult health: Turning gold into lead]. Z Psychosom Med Psychother 48: 359–369, 2002.

80. Felitti VJ. Adverse childhood experiences and adult health. Acad Pediatr 9: 131–132, 2009.

81. de Boo HA, and Harding JE. The developmental origins of adult disease (Barker) hypothesis. Aust N Z J Obstet Gynaecol 46: 4–14, 2006.

82. Louet JF, LeMay C, and Mauvais-Jarvis F. Antidiabetic actions of estrogen: insight from human and genetic mouse models. Curr Atheroscler Rep 6: 180–185, 2004.

83. Mauvais-Jarvis F. Estrogen and androgen receptors: regulators of fuel homeostasis and emerging targets for diabetes and obesity. Trends Endocrinol Metab 22: 24–33, 2011.

84. Vanitallie TB. Stress: a risk factor for serious illness. Metabolism 51: 40–45, 2002.

85. Ballestri S, Nascimbeni F, Baldelli E, Marrazzo A, Romagnoli D, and Lonardo A. NAFLD as a Sexual Dimorphic Disease: Role of Gender and Reproductive Status in the Development and Progression of Nonalcoholic Fatty Liver Disease and Inherent Cardiovascular Risk. Adv Ther 34: 1291–1326, 2017.

86. Fuller KNZ, McCoin CS, Von Schulze AT, Houchen CJ, Choi MA, and Thyfault JP. Estradiol treatment or modest exercise improves hepatic health and mitochondrial outcomes in female mice following ovariectomy. Am J Physiol Endocrinol Metab 320: E1020–E1031, 2021.

87. Curtis JR, Westfall AO, Allison J, Bijlsma JW, Freeman A, George V, Kovac SH, Spettell CM, and Saag KG. Population-based assessment of adverse events associated with long-term glucocorticoid use. Arthritis Rheum 55: 420–426, 2006.

88. Lofberg E, Gutierrez A, Wernerman J, Anderstam B, Mitch WE, Price SR, Bergstrom J, and Alvestrand A. Effects of high doses of glucocorticoids on free amino acids, ribosomes and protein turnover in human muscle. Eur J Clin Invest 32: 345–353, 2002.

89. Suter DE, and Schwartz NB. Effects of glucocorticoids on secretion of luteinizing hormone and follicle-stimulating hormone by female rat pituitary cells in vitro. Endocrinology 117: 849–854, 1985.

90. D’Agostino J, Valadka RJ, and Schwartz NB. Differential effects of in vitro glucocorticoids on luteinizing hormone and follicle-stimulating hormone secretion: dependence on sex of pituitary donor. Endocrinology 127: 891–899, 1990.

91. Schliep KC, Mumford SL, Vladutiu CJ, Ahrens KA, Perkins NJ, Sjaarda LA, Kissell KA, Prasad A, Wactawski-Wende J, and Schisterman EF. Perceived stress, reproductive hormones, and ovulatory function: a prospective cohort study. Epidemiology 26: 177–184, 2015.

92. Pal L, Bevilacqua K, and Santoro NF. Chronic psychosocial stressors are detrimental to ovarian reserve: a study of infertile women. Journal of psychosomatic obstetrics and gynaecology 31: 130–139, 2010.

93. Lapidus L, Bengtsson C, Larsson B, Pennert K, Rybo E, and Sjostrom L. Distribution of adipose tissue and risk of cardiovascular disease and death: a 12 year follow up of participants in the population study of women in Gothenburg, Sweden. Br Med J (Clin Res Ed) 289: 1257–1261, 1984.

94. Larsson B, Svardsudd K, Welin L, Wilhelmsen L, Bjorntorp P, and Tibblin G. Abdominal adipose tissue distribution, obesity, and risk of cardiovascular disease and death: 13 year follow up of participants in the study of men born in 1913. Br Med J (Clin Res Ed) 288: 1401–1404, 1984.

95. Valassi E, Scacchi M, and Cavagnini F. Neuroendocrine control of food intake. Nutr Metab Cardiovasc Dis 18: 158–168, 2008.

96. Considine RV, Sinha MK, Heiman ML, Kriauciunas A, Stephens TW, Nyce MR, Ohannesian JP, Marco CC, McKee LJ, Bauer TL, and et al. Serum immunoreactive-leptin concentrations in normal-weight and obese humans. N Engl J Med 334: 292–295, 1996.

97. Leon-Cabrera S, Solis-Lozano L, Suarez-Alvarez K, Gonzalez-Chavez A, Bejar YL, Robles-Diaz G, and Escobedo G. Hyperleptinemia is associated with parameters of low-grade systemic inflammation and metabolic dysfunction in obese human beings. Front Integr Neurosci 7: 62, 2013.

98. Agrawal S, Gollapudi S, Su H, and Gupta S. Leptin activates human B cells to secrete TNF-alpha, IL-6, and IL-10 via JAK2/STAT3 and p38MAPK/ERK1/2 signaling pathway. J Clin Immunol 31: 472–478, 2011.

99. Weisberg SP, McCann D, Desai M, Rosenbaum M, Leibel RL, and Ferrante AW. Obesity is associated with macrophage accumulation in adipose tissue. J Clin Invest 112: 1796–1808, 2003.

100. Munzberg H, Flier JS, and Bjorbaek C. Region-specific leptin resistance within the hypothalamus of diet-induced obese mice. Endocrinology 145: 4880–4889, 2004.

101. Enriori PJ, Evans AE, Sinnayah P, Jobst EE, Tonelli-Lemos L, Billes SK, Glavas MM, Grayson BE, Perello M, Nillni EA, Grove KL, and Cowley MA. Diet-induced obesity causes severe but reversible leptin resistance in arcuate melanocortin neurons. Cell Metab 5: 181–194, 2007.

102. Salzmann C, Otis M, Long H, Roberge C, Gallo-Payet N, and Walker CD. Inhibition of steroidogenic response to adrenocorticotropin by leptin: implications for the adrenal response to maternal separation in neonatal rats. Endocrinology 145: 1810–1822, 2004.

103. Schmidt MV, Levine S, Alam S, Harbich D, Sterlemann V, Ganea K, de Kloet ER, Holsboer F, and Muller MB. Metabolic signals modulate hypothalamic-pituitary-adrenal axis activation during maternal separation of the neonatal mouse. Journal of neuroendocrinology 18: 865–874, 2006.

104. Lopez M, Seoane LM, Tovar S, Garcia MC, Nogueiras R, Dieguez C, and Senaris RM. A possible role of neuropeptide Y, agouti-related protein and leptin receptor isoforms in hypothalamic programming by perinatal feeding in the rat. Diabetologia 48: 140–148, 2005.

105. Breton C, Lukaszewski MA, Risold PY, Enache M, Guillemot J, Riviere G, Delahaye F, Lesage J, Dutriez-Casteloot I, Laborie C, and Vieau D. Maternal prenatal undernutrition alters the response of POMC neurons to energy status variation in adult male rat offspring. American journal of physiology Endocrinology and metabolism 296: E462–472, 2009.

106. Delahaye F, Breton C, Risold PY, Enache M, Dutriez-Casteloot I, Laborie C, Lesage J, and Vieau D. Maternal perinatal undernutrition drastically reduces postnatal leptin surge and affects the development of arcuate nucleus proopiomelanocortin neurons in neonatal male rat pups. Endocrinology 149: 470–475, 2008.

107. Shin BC, Dai Y, Thamotharan M, Gibson LC, and Devaskar SU. Pre-and postnatal calorie restriction perturbs early hypothalamic neuropeptide and energy balance. Journal of neuroscience research 90: 1169–1182, 2012.

108. Baquero AF, de Solis AJ, Lindsley SR, Kirigiti MA, Smith MS, Cowley MA, Zeltser LM, and Grove KL. Developmental switch of leptin signaling in arcuate nucleus neurons. J Neurosci 34: 9982–9994, 2014.

109. Vaiserman AM. Early-Life Nutritional Programming of Type 2 Diabetes: Experimental and Quasi-Experimental Evidence. Nutrients 9: 2017.

110. Yam KY, Ruigrok SR, Ziko I, De Luca SN, Lucassen PJ, Spencer SJ, and Korosi A. Ghrelin and hypothalamic NPY/AgRP expression in mice are affected by chronic early-life stress exposure in a sex-specific manner. Psychoneuroendocrinology 86: 73–77, 2017.

111. de Lima RMS, Dos Santos Bento LV, di Marcello Valladao Lugon M, Barauna VG, Bittencourt AS, Dalmaz C, and de Vasconcellos Bittencourt APS. Early life stress and the programming of eating behavior and anxiety: Sex-specific relationships with serotonergic activity and hypothalamic neuropeptides. Behavioural brain research 379: 112399, 2020.

112. Eller-Smith OC, Nicol AL, and Christianson JA. Potential Mechanisms Underlying Centralized Pain and Emerging Therapeutic Interventions. Front Cell Neurosci 12: 35, 2018.

113. Friedman TC, Mastorakos G, Newman TD, Mullen NM, Horton EG, Costello R, Papadopoulos NM, and Chrousos GP. Carbohydrate and lipid metabolism in endogenous hypercortisolism: shared features with metabolic syndrome X and NIDDM. Endocrine journal 43: 645–655, 1996.

114. Beaupere C, Liboz A, Feve B, Blondeau B, and Guillemain G. Molecular Mechanisms of Glucocorticoid-Induced Insulin Resistance. Int J Mol Sci 22: 2021.

115. Desbriere R, Vuaroqueaux V, Achard V, Boullu-Ciocca S, Labuhn M, Dutour A, and Grino M. 11beta-hydroxysteroid dehydrogenase type 1 mRNA is increased in both visceral and subcutaneous adipose tissue of obese patients. Obesity 14: 794–798, 2006.

116. Atalar F, Gormez S, Caynak B, Akan G, Tanriverdi G, Bilgic-Gazioglu S, Gunay D, Duran C, Akpinar B, Ozbek U, Buyukdevrim AS, and Yazici Z. The role of mediastinal adipose tissue 11beta-hydroxysteroid d ehydrogenase type 1 and glucocorticoid expression in the development of coronary atherosclerosis in obese patients with ischemic heart disease. Cardiovasc Diabetol 11: 115, 2012.

117. Hageman RS, Wagener A, Hantschel C, Svenson KL, Churchill GA, and Brockmann GA. High-fat diet leads to tissue-specific changes reflecting risk factors for diseases in DBA/2J mice. Physiol Genomics 42: 55–66, 2010.

118. Morton NM, Ramage L, and Seckl JR. Down-regulation of adipose 11 beta-hydroxysteroid dehydrogenase type 1 by high-fat feeding in mice: a potential adaptive mechanism counteracting metabolic disease. Endocrinology 145: 2707–2712, 2004.

